# Closed microbial communities self-organize to persistently cycle carbon

**DOI:** 10.1101/2020.05.28.121848

**Authors:** Luis Miguel de Jesús Astacio, Kaumudi H. Prabhakara, Zeqian Li, Harry Mickalide, Seppe Kuehn

## Abstract

Nutrient cycling is an emergent property of ecosystems at all scales, from microbial communities to the entire biosphere. Understanding how nutrient cycles emerge from the collective metabolism of ecosystems is a challenging problem. Here we use closed microbial ecosystems (CES), hermetically sealed consortia that sustain nutrient cycles when provided with only light, to learn how microbial communities cycle carbon. A new technique for quantifying carbon exchange shows that CES comprised of an alga and diverse bacteria self-organize to robustly cycle carbon. Comparing a library of CES, we find that carbon cycling does not depend strongly on the taxonomy of the bacteria present. Metabolic profiling reveals functional redundancy across CES: despite strong taxonomic differences, self-organized CES exhibit a conserved set of metabolic capabilities.

**Summary:** Closed microbial communities of algae and bacteria self-organize to robustly cycle carbon via emergent metabolite exchange.

---

Nutrient cycles are a defining emergent property of ecosystems at all scales. Ecosystem persistence relies on nutrient cycles to continuously replenish resources. As a result, global cycles of carbon (*1*) and nitrogen (*2*) are key organizing processes of life across the planet. In microbial communities nutrient cycling is also an key functional process, from carbon fixation and respiration in microbial mats (*3*), to denitrification and nitrogen fixation in soils (*4*), sulphur cycling in anaerobic marine microbial communities (*5*), and nutrient cycling in periphytic consortia (*6*).

The fact that nutrient cycling is an essential feature of ecosystems means that a key problem in ecology is understanding how the cyclic flow of nutrients emerges from interactions between organisms in communities (*7*). Microbial communities, owing to their small size, rapid replication rates and tractability in the laboratory, are powerful model systems for discovering the principles governing ecosystem organization and function. For example, a conserved succession of bacteria with predictable metabolic capabilities describes the degradation of particulate organic carbon in marine microbial communities (*8*). Complex bacterial communities propagated in the laboratory reveal emergent cross-feeding between predictable taxa (*9*), and simple assembly rules govern the stable composition of synthetic communities (*10*).

However, few quantitative studies have exploited the advantages of microbial communities in the laboratory to uncover the principles governing nutrient cycling. A primary roadblock to studying nutrient cycling in model microbial communities is experimental: most existing approaches use batch (*9*) or continuous culture (*11*), where nutrients are supplied externally. In these conditions, with few exceptions (*12, 13*), nutrient cycling rarely occurs since the external supply of nutrients favors those strains that can most rapidly exploit the supplied resource (*8,9*). The continuous dilution of these systems means that slower growing taxa are quickly washed out of the system (*14*), frequently resulting in the assembly of communities with taxa that either exploit the primary resource or are sustained via strong mutualistic or commensal interactions (*9, 15*). In contrast, nutrient cycling means that not all nutrients are supplied exogenously, but instead that some nutrients are regenerated by the community itself. Stable nutrient cycling therefore requires a balance between the production of byproducts (e.g. CO_2_ by respiration) and their consumption (CO_2_ fixation by photosynthesis) in a closed loop. Here we seek to develop microbial communities as model systems to understand how communities are organized to cycle nutrients.

To address this problem, we built on the work of Folsome (*16*), Taub (*17*) and others to develop closed microbial ecosystems (CES) as models for understanding the principles of emergent nutrient cycling. CES are milliliter-scale aquatic communities which are hermetically sealed and illuminated (*16–20*). Since no nutrients enter or leave a CES after assembly, persistence in these communities requires that nutrient cycles be sustained through photosynthesis. Complex CES have been shown to retain biological activity for decades in some cases (*20*). As such, CES are ideal model microbial ecosystems for understanding nutrient cycling (*21*). However, most work on CES to date has focused on applications to spaceflight (*22*) or population dynamics (*19, 23, 24*) rather that understanding the emergent organization of ecosystems that cycle nutrients.

Here we take a top-down approach (*9, 16*) to assemble a library of CES, comprising diverse bacterial consortia and a single algal species. We present a new, high-precision, method for quantifying carbon cycling *in situ* to show that our CES rapidly and persistently cycle carbon. We utilize sequencing and metabolic profiling to reveal the conserved features of CES that cycle carbon.

Carbon cycling arises in a CES from the complementary reactions of photosynthesis and respiration, which consume (produce) and produce (consume) CO_2_ and O_2_, respectively (Figure 1A). Carbon cycling emerges from the photosynthetic conversion of CO_2_ into organic carbon which is then either excreted by phototrophic microbes (*25*) or made available to bacterial decomposers via death of primary producers. The subsequent respiration of organic carbon by bacterial community members produces CO_2_, completing the cycle.

**Figure 1:**
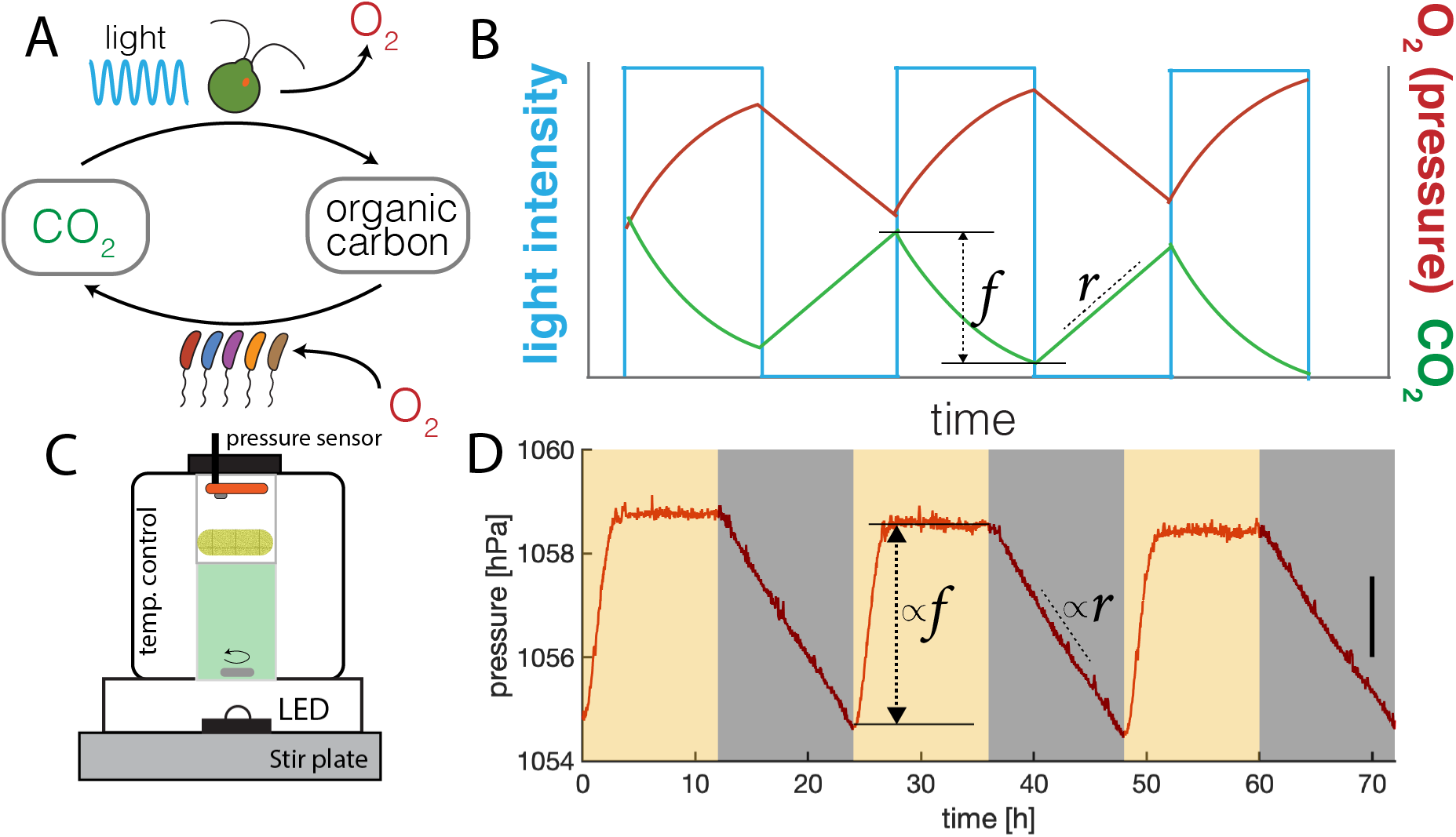
Quantifying carbon cycling in closed microbial ecosystems. (A) Schematic of carbon cycling in closed ecosystems which occurs via photosynthesis utilizing light to fix CO_2_ to organic carbon and produce O_2_ (top arrow) and respiration which utilizes O_2_ and organic carbon to produce CO_2_. (B) Sketch of changes in total O_2_ or pressure (red line) and CO_2_ (green line) in a CES subjected to cycles of light and dark (blue line). Sketch assumes photosynthetic rate exceeds respiration rate during the light phase. *r* is the rate of increase of CO_2_ during the dark phase. *f* is the net decrease in CO_2_ during the light phase. Assuming respiratory and photosynthetic quotients of one, O_2_ dynamics mirror CO_2_. Since O_2_ is 30-fold less soluble in water than CO_2_ changes in pressure quantify changes in O_2_ and CO_2_ concentrations in a CES (Supplementary Appendix). (C) A schematic of our custom cultivation devices for quantifying carbon cycling in CES using pressure sensors. 20 mL CES are housed in glass vials (40 mL total volume), stirred at 450 rpm, illuminated by an LED and held at 30 *°*C under feedback temperature control (Supplementary Appendix). A high-precision pressure sensor is integrated into the hermetically sealed cap and a porous foam stopper (yellow) shades the sensor from illumination. (D) Pressure measurements (acquired once per second) in a CES subjected to 12 h-12 h light-dark cycles as indicated by orange and gray shaded regions respectively. Light intensity during the light phase is 150 µmol m^*−*2^ s^*−*1^. Pressure rises and falls in response to light and dark as expected. The pressure stabilizes during the light phase, indicating that photosynthesis becomes CO_2_-limited. The change in pressure is proportional to *r* and *f* as labeled. Carbon cycling, computed from these quantities, is proportional to the amplitude of pressure oscillations (Supplementary Appendix). Data in (D) are smoothed with a one minute moving average. A change in pressure of 1.56 hPa (black line, right side) corresponds to a production/consumption of approximately 2 µmol of CO_2_ assuming pH 6.5 and photosynthetic/respirtory quotients of 1. (Supplementary Appendix).

Carbon cycling can be quantified by continuously measuring O_2_ or CO_2_ production and consumption in a CES subjected to cycles of light and dark (*17*). The dependence of photosynthetic O_2_ production (CO_2_ fixation) on light means that diel cycles of light and dark result in oscillations in O_2_ and CO_2_ levels (Figure 1B). To estimate carbon cycling rates from measurements of O_2_ or CO_2_ dynamics, one measures the rate of respiration during the dark phase (*r*, Figure 1B,D). We then assume that this rate is sustained during the light period, allowing us to compute the total CO_2_ produced during a light-dark cycle. The amount of CO_2_ fixed during the light phase can then be computed by measuring the net oxygen production (CO_2_ fixed, *f*, Figure 1B) during the light phase and using the respiration rate to infer a total CO_2_ fixed (Supplementary Appendix). The quantity of carbon cycled over a light-dark cycle is then the number of moles of inorganic carbon both fixed and produced. Assuming fixed photosynthetic and repiratory quotients (ratio of O_2_ production (consumption) to CO_2_ consumption (production)) allows carbon cycling to be quantified by measuring either O_2_ or CO_2_ dynamics. Obenhuber and Folsome have shown that the 30-fold lower solubility of O_2_ relative to CO_2_ in water results in oscillations in pressure in a sealed vessel which are proportional to changes in O_2_ and therefore CO_2_ in the community (*16*). Similar methods are used to measure primary production in aquatic ecosystems in the wild (*26*).

We developed a custom culture device to precisely measure changes in pressure in a CES subjected to cycles of light and dark. A schematic is shown in Figure 1C. Each device housed a 20 mL CES in a 40 mL glass vial. The cap of the hermetically sealed vial was fitted with a high-precision low-cost, pressure sensor developed for mobile devices (Bosch, BME280). In contrast to direct detection of O_2_ or CO_2_, pressure measurements are higher sensitivity, lower cost, require no calibration, do not consume analyte and are stable for months. The vial was illuminated from below by a light-emitting diode (LED) and fit in a metal block which was held under feedback temperature control via a thermoelectric heating-cooling element (*27*). When we subjected the CES housed in our devices to cycles of light and dark (12 h-12 h), we observed increases and decreases in pressure, as expected (Figure 1D). The respiration rate (*r*) and net productivity *f* can be quantified directly from these continuous pressure measurements. The rate of carbon cycling in our CES is proportional to the amplitude of the light-driven pressure oscillations (Supplementary Appendix). Performing the same experiment with only water in the vial resulted in no pressure oscillations as expected (Figure S1), and concurrent measurements of O_2_ and pressure in the vial confirmed that pressure changes reflected the production and consumption of O_2_ and therefore CO_2_ (Figures S2-S3).

Using these devices, we initially measured carbon cycling in variants of a previously studied synthetic CES (*23, 24*) comprised of *Chlamydomonas reinhardtii* (UTEX 2244, mt+) and *Escherichia coli* (MG1655) over periods of several weeks. We found that these simple synthetic communities failed to persistently cycle carbon (Figure 2C, Figure S4). We speculate that this failure arose from the production of starch by the algae (*28*) which cannot be utilized by *E. coli*. We reasoned that increasing the metabolic diversity of the bacterial component of our CES might improve carbon cycling. To accomplish this, we turned to a top-down community assembly approach (*9, 11*) outlined in Figure 2A.

**Figure 2:**
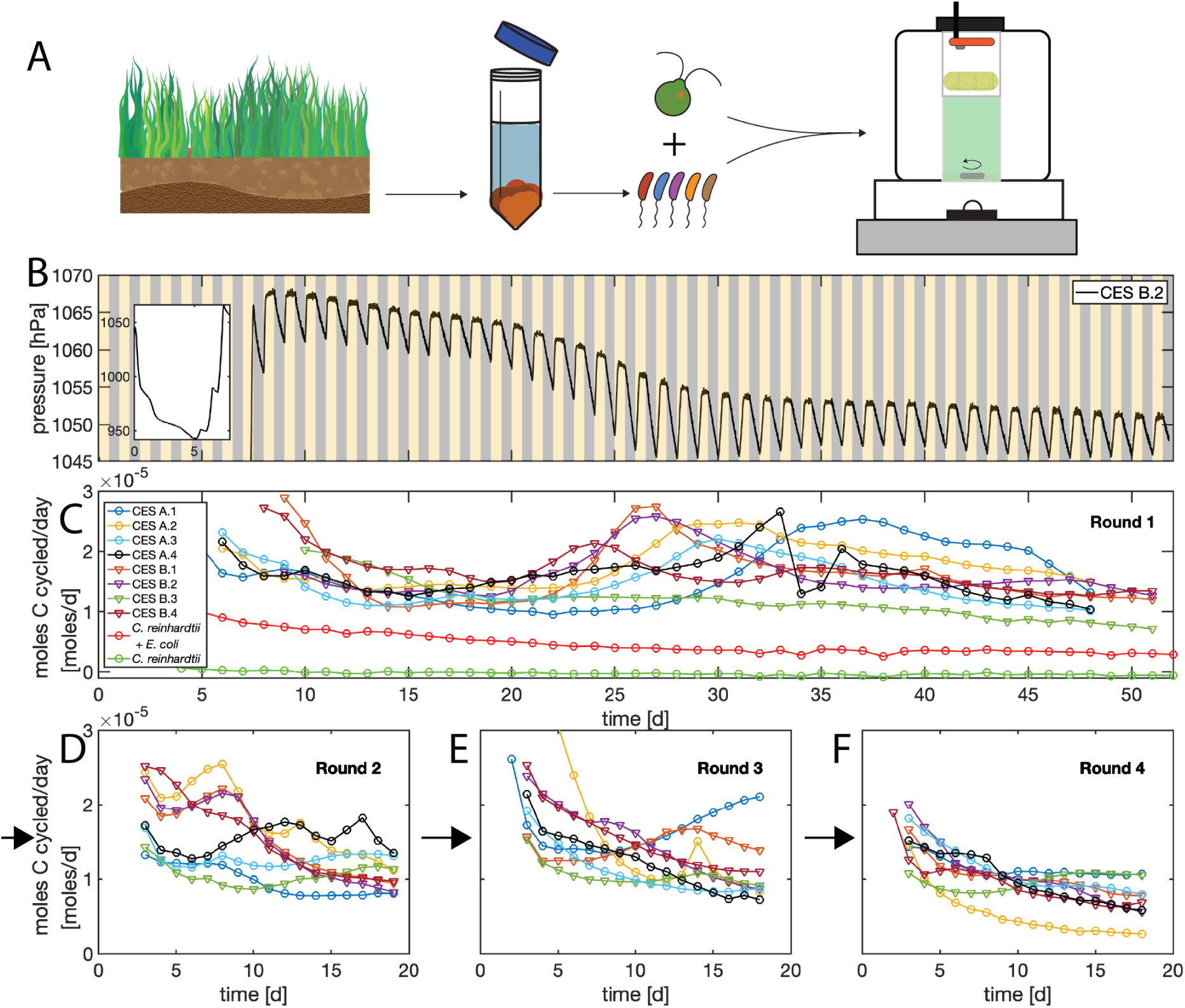
Long-term carbon cycling in closed ecosystems comprised of *Chlamydomonas reinhardtii* and soil-derived bacterial communities. (A) Top-down assembly of microbial CES. Soil samples are harvested and bacterial communities are extracted. Bacteria are then combined with the alga *C. reinhardtii* and inoculated into the custom culture devices described in Figure 1C. Eight CES were assembled, four each from two soil samples (“A” and “B”) in defined minimal medium, and subjected to 12 h-12 h light-dark cycles (orange/gray shaded regions) for *∼*50 days while pressure was measured. Light intensity was 150 µmol m^*−*2^ s^*−*1^ during light phase. (B) Pressure measurements performed once per second, smoothed by a one minute moving average, for one of the eight CES. The initial large drop in pressure due to rapid respiration of supplied organic carbon (glucose) is shown in the inset. The rate of carbon cycling (moles/day) for all eight CES is computed from pressure traces as described in the Supplementary Appendix. Carbon cycling rates are only reported after the initial transient phase (inset, panel A) has ended. We assume respiratory and photosynthetic quotients of 1, and a pH of 6.5. Circles indicate CES from soil sample A and triangles from sample B. The transient increase in cycling around 25 to 35 days coincides with a reduction in photosynthetic rates and an increase in respiration (Figure S7). Red and green traces are synthetic CES comprised of *C. reinhardtii E. coli* (mean of two replicates) and *C. reinhardtii* (single replicate, Figure S4) as shown in the legend. Statistical errors in estimates of carbon cycling are smaller than the size of the markers. Legend in (C) applied to (D-F). At the end of the acquisition shown in (C) all eight CES were opened, samples were taken and CES were diluted 1:20 into fresh media. CES were then sealed for an additional *∼* 18 d of light-dark cycles and carbon cycling was monitored. (D) Shows carbon cycling rates after the first dilution. Two additional dilution rounds were performed and cycling rates are shown in (E-F) as indicated by the black arrows. The average cycling rates at the end of each round do not differ significantly between rounds of enrichment (p-values: 0.31,0.87 and 0.053, two-sample t-test between last measurement between rounds 1 and 2, 2 and 3, 3 and 4 respectively)

To assemble communities, we sampled local soils, removed eukaryotes by applying drugs, and extracted bacterial communities using standard techniques (Supplementary Appendix). We then combined these diverse bacterial populations with the domesticated soil dwelling alga *C. reinhardtii* (Figure 2A). The resulting CES contained a diverse assemblage of bacteria and a single, well characterized, photosynthetic microbial species. We assembled 8 CES using this method, 4 each from two soil samples (designated “A” and “B”), and inoculated them into a chemically defined freshwater mimic medium (*29*) which included organic carbon (glucose), nitrogen (ammonia) and phosphorous (phosphate, Table S4) to facilitate the initial growth of the community. We then sealed these communities in vials and placed them in culture devices like the one shown in Figure 1C and subjected them to 12 h-12 h light-dark cycles for a period of approximately 50 d.

A representative time series of pressure for one of these CES is shown in Figure 2B. We observed an initial large decline in pressure (Figure 2B, inset) which arose from the rapid bacterial respiration of glucose (this decline is not present in CES of algae alone, Figure S4). The pressure remains approximately 10 % below ambient for 5 to 8 days and then begins to rise (Figure S5), reflecting the timescale over which we expect algae to grow (*30*). The rising pressure reflects photosynthetic activity (O_2_ production) by the alga before saturating after 8 to 10 days (Figures S2, S5). Once the pressure saturated, we observed stable oscillations in the pressure driven by light-dark cycles. In this regime, during each light phase, the pressure stabilized within 2 to 3 hours of the illumination being turned on. Therefore, the algae rapidly fix CO_2_ early in the light phase before exhausting the inorganic carbon supply later in the light phase. After CO_2_ is depleted during the early periods of the light phase, respiration and photosynthesis are balanced resulting in stable pressure (O_2_ levels) late in the light phase. We conclude that the respiration is the rate-limiting step in the carbon cycle in our CES, and that light is not limiting algal carbon fixation. During the dark phases of each light-dark cycle, we observe a linear decrease in pressure with time, indicating a constant rate of respiration during the dark phase (Figure S6).

We observed stable pressure oscillations, with saturating pressure levels during the light phase and constant respiration rates during the dark phase, for a period of approximately 50 d. During this period, we observe longer timescale dynamics whereby the pressure (O_2_) levels slowly drop after about 25 d (7/8 CES, Figure 2B, Figure S5). A detailed analysis of the O_2_ dynamics reveals that this decline in pressure coincides with a slowing of the photosynthetic rates and an increase in the respiration rates (Figure S7). We hypothesize that this results from the death of a fraction of the algal population which supplies the bacterial community with additional organic carbon for respiration.

We estimated the rate of carbon cycling in each of our 8 CES directly from pressure measurements like the one shown in Figure 2B and the results are shown in Figure 2C. We observe robust carbon cycling at rates of approximately 10 to nearly 30 *µ*mol d^*−*1^ in all 8 CES. The magnitude of this carbon cycling rate is a sizable fraction of the total organic carbon supplied to each CES at the outset (*∼*200 *µ*mol, Table S5), and the amount of non-volatile organic carbon present in each CES at the end of the experiment (120 *µ*mol to 180 *µ*mol, Figure S8). Therefore, in a period of between 4 and 20 days the amount of carbon cycled approaches the total carbon in the CES. In this sense, we conclude that the carbon cycling rate in our self-assembled CES is high. In contrast, in CES comprised of *C. reinhardtii* or *C. reinhardtii* and *E. coli* we observe carbon cycling rates that are below our detection limit, and *∼*4-fold lower than the complex CES, respectively (Figure 2C, green and red circles). We conclude that CES comprised of *C. reinhardtii* and complex soil-derived bacterial communities self-organize to rapidly cycle carbon.

How do similar carbon cycling rates across CES emerge from bacterial consortia derived from distinct soil samples? One possibility is taxonomic similarity between assembled bacterial communities. In this scenario, one or a few similar bacterial taxa would rise to high abundance potentially due to their ability to utilize the organic carbon produced by *C. reinhardtii* (*25*). Another possibility is that taxonomically distinct consortia are maintained in each CES despite the similar carbon cycling rates, and it is the metabolic capabilities of the assembled bacterial communities that is similar from one CES to the next and not the taxa present. The latter outcome could arise from functionally redundant bacterial communities (*11,31*) that are able to consume the available organic carbon but are comprised of taxonomically distinct bacteria.

To test between these possibilities we performed an enrichment experiment that allowed us to quantify the taxonomic composition and metabolic properties of our CES, while enriching communities for those taxa essential for carbon cycling. Each CES was opened, sampled, assayed and diluted 1:20 into fresh medium over three rounds. We chose three rounds of 1:20 dilution to reduce the abundance of any strains not able to grow in our CES by 8000-fold, putting them below our detection limit by sequencing. Each CES was opened after an initial 50 d period of closure (round 1), and diluted before being sealed to continue the carbon cycling measurement. At each round, samples were taken from each CES for metabolic assays and sequencing. The enrichment was performed three times (rounds 2-4) with *∼*18 day periods of closure for each round. Carbon cycling rates during each of these enrichment phases are shown in Figure 2D-E (Figure S9). In most CES, we observed a decline in carbon cycling rates during the first approximately 10 days of closure before rates stabilize across most CES. We found that the average cycling rates at the end of each round of dilution do not differ significantly from one round to the next (Figure 2). However, one of eight CES exhibited a substantial decline in carbon cycling rates relative to the mean in the final round (CES A.2, yellow curve, Figure 2F). Further, two CES were diluted and sealed again after round 4 and showed stable cycling for an additional period of *>*130 days (Figure S10). We conclude that the carbon cycling is robust to serial dilution and that our CES can stably cycle carbon for many months.

Between each of the four rounds of enrichment (Figure 2C-F) samples were taken from each of the 8 CES. On each of these samples, we performed 16S amplicon sequencing (V4 hypervariable region) of bacterial communities. Figure 3A shows a time series of the dominant taxa in all 8 CES across all four rounds of dilution (dominant taxa are those at relative abundance above 5 % in any time point). We find that the bacterial communities in our CES differ strongly from the initial soil samples (Figure S11), indicating that closure and the presence of algae results in a dramatic re-organization of the soil community. Taxonomically, assembled CES comprise *>*5 taxa which make up approximately three quarters of the population in each community. Some CES exhibit relatively large taxonomic variation from round to round (A.1, A.2 and A.4), while in others we find that the taxonomic structure of each CES is relatively stable from round to round (A.2, B.2, B.3, Figure 3A). While all CES from soil sample B harbor a taxon from the genus *Terrimonas*, the same taxon is only observed in later round of enrichment in one of the four CES from sample A. Further, all CES retain between 80 and 220 rare taxa (relative abundances *<*5 %) with the number declining after round 1 (Figure S12). Therefore, a visual inspection of Figure 3A suggests that there is no obviously conserved taxonomic structure across our CES. To better quantify this observation, we computed the Jensen-Shannon divergence (JSD) (*32*) between the relative abundances in each pair of CES at each round of enrichment. The JSD quantifies differences in community composition between two communities and varies between 0 for two identical communities and 1 for two communities that share no taxa in common. On average, the taxonomic composition differs more between CES (inter-CES) than it does for the same CES across rounds of enrichment (intra-CES, Figure S13), a result that is robust to using other community similarity metrics (Figure S14). We also found that the JSD between CES from different soil samples did not decline across rounds of enrichment (Figure S15), indicating that the taxonomic differences between CES from different soil samples are retained through the enrichment process. Inter-CES divergences remained larger than intra-CES divergences even when we grouped taxa with only 90 % 16S sequence similarity, indicating that there is not taxonomic similarity between CES even at higher levels of classification (Figure S16, S17). To visualize community taxonomic composition, we embedded the JSD between all CES at all rounds into two dimensions using multi-dimensional scaling (MDS) (Figure S18 quantifies the stress of this embedding) and the result is shown in Figure 3B. Note that the CES remain largely separated from each other in this embedding. Figure 3B supports our assertion that the taxonomic composition differs strongly from one CES to the next and that during enrichment these differences are retained. The differences between CES from soil sample A are larger than those for sample B (Figure S13), but in neither case did we observe CES converging to a shared taxonomic makeup of the bacterial community. We conclude that the bacterial communities in our CES differ substantially in their taxonomic composition.

**Figure 3:**
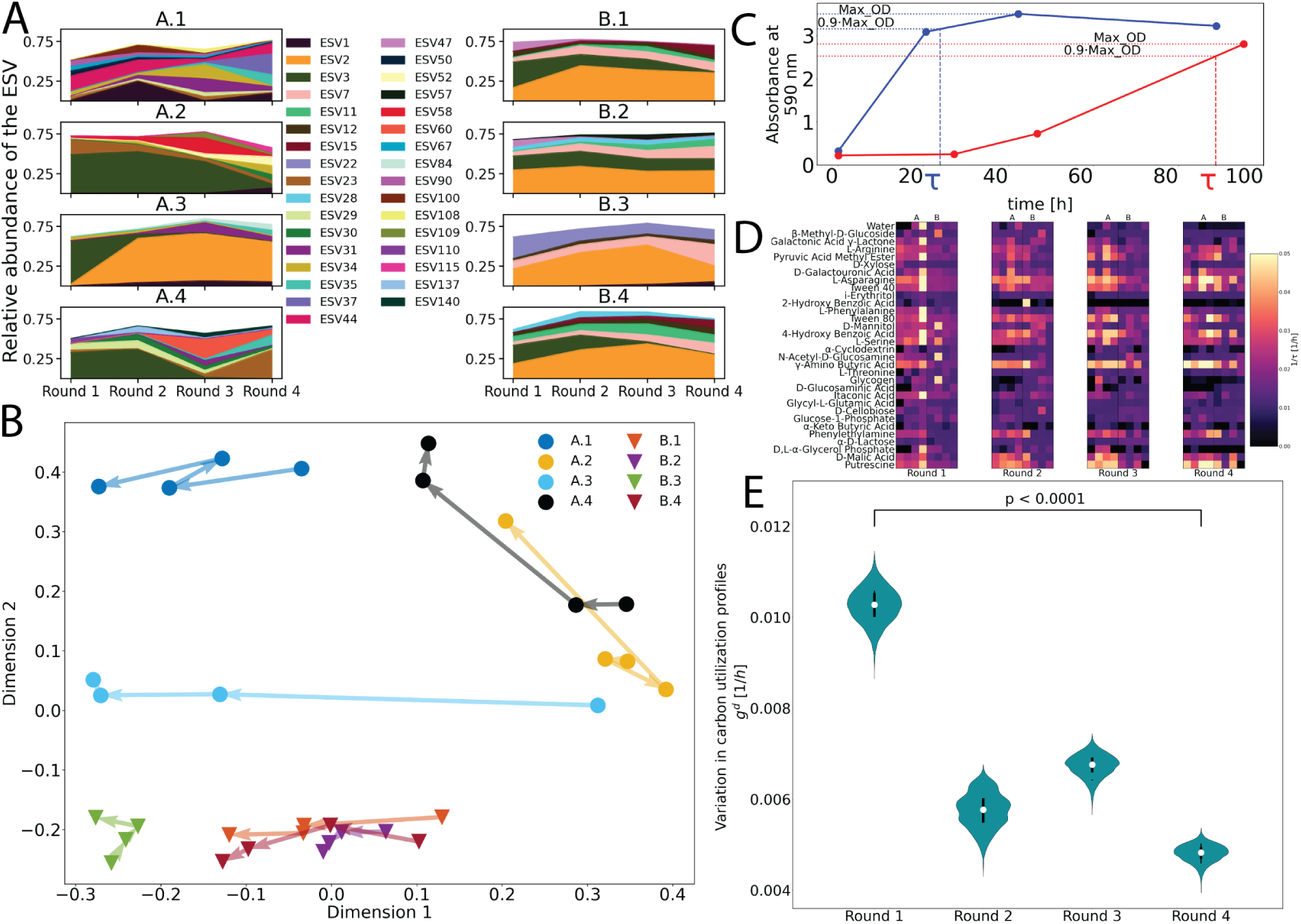
Divergent taxonomic structure and convergent metabolic capabilities across replicate CES. (A) Relative abundances measured by 16S rRNA amplicon sequencing of the bacterial taxa comprising the CES (y-axis) for each round of dilution (x-axis). Each exact sequence variant (ESV) is represented by a unique color, indicated in the legend. Only the ESVs that have a relative abundance of 5 % or higher in at least one of the four dilution rounds for each CES are shown. Most ESVs belong to unique genera (Figure S24, where multiple ESVs having the same genus are combined). (B) The Jensen Shannon Divergence (JSD) of the relative abundances of all detected taxa at the ESV level is computed between all the 32 CES, as described in the Supplementary Appendix. Multi-dimensional scaling (MDS) is applied to the JSD to embed the data in two dimensions. The circles denote CES derived from soil sample A and the triangles denote CES derived from soil sample B, colors correspond to Figure 2C. The arrows indicate transitions between dilution rounds. (C) Two time series of absorbance (590 nm) indicating respiration in Biolog EcoPlates via the redox sensitive dye tetrazolium (*33*). For each time series we compute a timescale (*τ*) by finding the the maximum absorbance (Max OD in the figure). We then linearly interpolate between measurements to compute *τ* as the time to reach 0.9*×*Max OD. In the two example measurements shown here, the blue curve reaches its maximum absorbance faster (*τ* = 23.96 h), indicating more rapid carbon uptake, while the red reaches it slower (*τ* = 88.79 h), indicating a slower utilization of the carbon source. For time series that do not show an increase in OD_590_ of at least 0.3 we assume no respiration and set *τ→* ∞ (Supplementary Appendix). In panels (D-E) we consider the quantity 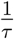. After each dilution round, we measured 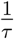 for 32 carbon sources, each in triplicate. The carbon respiration profiles of the eight CES are shown here for each dilution round, with carbon sources in rows and CES in columns. Dilution rounds are shown in separate panels (left to right) as labeled below. In each panel, CES from soil sample A are shown on the left and B on the right. Each entry indicates a mean 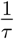 across the three replicate measurements for each carbon source in each CES. Lighter colors indicate faster consumption (smaller *τ*) of the carbon source. (E) Shows the decline in variability of carbon utilization profiles from rounds 1 to 4. The geometric mean variability in carbon utilization rates is computed as follows. Let 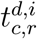 be the consumption rate of carbon source *i* for the *r*^*th*^ replicate of the *c*^*th*^ CES at dilution round *d* (*r* ∈{1, 2, 3}, *c* ∈{1, 2, 3, *…*, 8},*i* ∈{1, 2, 3, *…*, 32}, and *d* ∈{1, 2, 3, 4}). For each carbon source at each dilution round we compute *σ*^*d,i*^(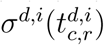), which is the standard deviation in the carbon utilization rate across all CES (*c*) and replicate measurements (*r*) for each carbon source in a given dilution round. An aggregated measure of variability in carbon utilization rates for each dilution round *d* is obtained by computing the geometric mean of *σ*^*d,i*^ across carbon sources: 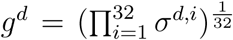. This quantity is plotted for each round of enrichment. Errors in *g* were computed by bootstrap re-sampling each 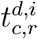 (across *r*) 10,000 times to generate 10,000 resampled values of *g*^*d*^. To test for significance we compute the difference in the geometric mean between dilution rounds 1 and 4 for each bootstrapped replicate and computed the fraction of differences below zero.

The result that the taxonomic structure differs strongly from one CES to the next despite similar carbon cycling rates supports the idea that carbon cycling in our CES is accomplished by diverse but functionally redundant bacterial communities. In this case, we hypothesized that the metabolic capabilities of the assembled bacterial communities might be conserved across CES. Reasoning that the identity of the organic carbon compounds produced by *C. reinhardtii* is likely similar across CES, we hypothesized that the carbon utilization capabilities of the assembled bacterial communities might be similar across CES.

To test this hypothesis we measured carbon utilization capabilities on a diverse library of carbon sources for all CES after each round of enrichment. To accomplish this we used Biolog 96-well EcoPlates (*33*) which exploit a redox sensitive dye to report respiration in the presence of 32 diverse carbon sources (including compounds excreted by *C. reinhardtii*, Table S6) each in triplicate. After each round of dilution we distributed aliquots of each CES into an EcoPlate. We then incubated the plates and measured dye absorbance, a proxy for carbon utilization, daily for a period of 4 days. Example absorbance time series are shown in Figure 3C. For each replicate of each carbon source, we computed a timescale of respiration for that carbon source (*τ*). To compute *τ* we took the maximum absorbance detected over the course of the time series, and then computed the time to reach 90 % of that maximum (dashed lines, Figure 3C). The quantity 1*/τ* quantifies the rate at which a CES utilizes a given carbon compound. We averaged 1*/τ* across the three replicates for each carbon source at each round of enrichment in each CES (Figure 3D). Each row of Figure 3D shows the average 1*/τ* (utilization rate) for a single carbon source and each column a profile for a CES. Comparing carbon utilization profiles across rounds reveals a convergence in the metabolic capabilities across our 8 CES, with profiles becoming more similar across CES as the number of rounds of dilution increases. For example, by the end of round 4 none of the CES utilize 2-hydroxy benzoic acid despite 6 of 8 CES being capable of consuming the carbon source after round 1. Conversely, the enrichment process increases 1*/τ* for other carbon sources (phenylethylamine, putrecine, *γ*-amino butyric acid). We note that the carbon utilization profiles of the enriched CES, after round 4, differ strongly from *E. coli* (Figure S19) which itself fails to cycle carbon with *C. reinhardtii* (Figure 2C), suggesting that the carbon utilization capabilities of the complex CES are important for stable carbon cycling.

To quantify the variation in the carbon utilization profiles (columns, Figure 3E) across CES we computed the standard deviation in the rate of carbon utilization (1*/τ*) for each carbon source across all CES in each round of enrichment (*σ*^*i*^, where *i* indexes carbon sources). Large values of this standard deviation indicate large differences in carbon utilization rates across CES, and small values of this standard deviation indicate similar utilization rates for a given carbon compound. On average, across all 32 carbon compounds, we observe a decline in the standard deviation from round 1 to 4 (Figure S20), indicating that the carbon utilization profiles become more similar across CES. To better visualize this convergence across CES in carbon utilization profiles, we computed the geometric mean of the *σ*^*i*^ across all carbon compounds for each round of enrichment. The geometric mean was used since it captures the fractional change in variation across CES in the utilization rates of each carbon compound. Using this metric, we observed a substantial decline in the variability in carbon utilization rates across CES from rounds 1 and 4 (Figure 3E). We conclude that the CES are converging to similar carbon utilization profiles.

The fact that our CES exhibit similar carbon utilization profiles and carbon cycling rates suggests that these CES have been assembled under carbon limitation. Our pressure data show that photosynthesis by *C. reinhardtii* is CO_2_ limited and our media was designed with nitrogen and phosphorous in excess (Tables S4 and S5). We speculated that the metabolic convergence we observe in Figure 3E might be a consequence of carbon limitation in our CES, forcing the bacterial community to consume specific sets of carbon compounds produced by the algae. Indeed a control experiment indicates that some of the compounds utilized by the assembled bacterial communites are excreted by *C. reinhardtii* (Table S6, Supplementary Data 3). However, from the pressure data or metabolic profiling, we cannot determine the nutrient limiting respiration in our CES. To address this question we performed an assay after each round of dilution to determine the nutrient limiting respiration. We used a Microresp assay (Supplementary Appendix) whereby small aliquots of each CES were dispensed into 96-well plates and supplemented with carbon, nitrogen or phosphorous. We measured CO_2_ production in each sealed well directly using a pH sensitive dye and compared the results to control wells where no nutrients were added (Figure S21). We found respiration in our CES was in some cases carbon limited, but in many instances was phosphorous limited (predominantly in CES from soil sample A). In one CES, the identity of the limiting nutrient changed from one round to the next (CES A.2, Figure S21). Therefore, the metabolic convergence we observe across CES arises despite the fact that respiration is not limited by carbon in all CES. A quantitative analysis of the nutrient budgets in our CES revealed that phosphorous limitation must arise from phosphate accumulation, either by bacteria (*34*) or *C. reinhardtii* (*35*) and not the incorporation of phosphorous into biomass (Supplementary Appendix). We conclude that the self-organized carbon cycle in our CES, and the convergent carbon source utilization repertoire, is robust to changes in the identity of the nutrient limiting respiration.

Our results support the idea that carbon cycling in microbial communities can be sustained with functionally redundant bacterial consortia that exhibit a conserved set of metabolic capabilities despite variability in their taxonomic structure. Moreover, carbon cycling appears to be robust to differences in the identity of the nutrient limiting respiration in the CES. The result points to the idea that the emergent functional property of carbon cycling in microbial ecosystems is likely to arise from a conserved set of metabolic capabilities (*31*), that is robust to variation in taxonomic and nutrient limitation variation of the system. Our data suggest that the conserved properties of carbon cycling CES are likely carbon utilization pathways and the taxonomic diversity in our CES potentially reflects the weak phylogenetic conservation of carbon utilization phenotypes (*36*).

We have established CES as model systems for understanding how nutrient cycles emerge from metabolic processes in microbial communities. We propose that the CES studied here constitute powerful model systems for the detailed study of emergent nutrient cycling in ecosystems. For example, it will be interesting to extend this study to understand how this taxonomic variability and metabolic convergence impacts the stability of nutrient cycling. Quantifying abundance dynamics and metabolite exchanges *in situ* should reveal how interactions endow these communities with stable cycling capabilities, and permit comparing our experiments to theoretical work on closed ecologies (*37–40*). Further, the essential exchange of metabolites between photosynthetic and heterotrophic organisms in our CES means these systems can be used to study the role of co-evolution in ecosystem function.

## Acknowledgments

We acknowledge Dr. Karna Gowda for assistance with the microresp assay, James O’Dwyer and Andrew Ferguson for useful discussions, and Annette Wells for laboratory support. The Raymond J. Carver Biotechnology Center at the University of Illinois at Urbana-Champaign. LMJA and ZL acknowledge support from The Center for the Physics of Living Cells graduate fellowship program (National Science Foundation, PHY 0822613 and PHY 1430124).

## Supplementary materials

Supplementary Appendix

Figs. S1 to S24

Tables S1 to S6

References *(41-72)*

