## Supplementary Information for "Closed microbial communities self-organize to persistently cycle carbon"

### 1 Collecting and processing soil samples

The soil samples were collected on October 22, 2018 around 14:00 CT from two locations about 100 m apart from a restored prairie (Meadowbrook Park, Urbana, IL) located at 40°04'42.9"N and 88°12'22.3"W. The soil was dug to a depth of about 5 cm using autoclaved steel scoopula. Soil was collected from the bottom of the hole to minimize the probability of collecting native photosynthetic bacteria. The collected soil was placed in sterile 50 mL Falcon tubes. Fresh gloves and scoopula were used for each dig to minimize cross contamination.

About 5 g of soil was transferred to 15 mL Falcon tubes and about 10 mL MilliQ water was added to each tube. The tubes were strongly vortexed for about a minute. The soil was sufficiently soft for the vortexing to break down the particles. The soil was allowed to settle for 25 min. A small volume of the supernatant was used to measure pH using a pH paper. For both soil samples, the pH was between 6 and 6.4. The supernatant in the Falcon tubes was transferred to Eppendorf tubes and centrifuged at 7000 rpm for 5 min. The Falcon tubes with the rest of the soil were stored at 4 C. The supernatant was discarded and the pellet was re-suspended in an equal volume of the experimental media. The drugs cycloheximide (SKU - C7698 from SigmaAldrich) and nystatin (SKU - N4014 from SigmaAldrich) were added at concentrations of 200 $\mu$g/mL and 20 mg/L respectively. Cycloheximide inhibits protein synthesis in eukaryotic cells and is used here to terminate any eukaryotes present in the soil sample. Nystatin is used as a fungicide to target any fungi present in the soil samples. The samples are placed in sterile test tubes wrapped in aluminium foil. These test tubes were shaken at 225 rpm at 30 °C in an orbital shaker for 48 h. The aluminium foil blocks light, thus preventing the growth of any obligate photoautotrophs.

After 48 h, 1mL aliquots of the samples are transferred to sterile Eppendorf tubes and centrifuged at 7000 rpm for 7 minutes. The supernatant is discarded and the pellet re-suspended in fresh experimental media of equal volume. The same washing procedure is repeated once more (two washes in total). Washing removes the drugs, so that the growth of *Chlamydomonas reinhardtii*, a eukaryote added in the subsequent steps, is not inhibited. The contents of the Eppendorf tubes were then combined into a single Falcon tube for each soil sample and used to initiate CES as described below. The recollection of same-soil material into a single Falcon tube is done to guarantee homogeneity of initial community structure for all CES inoculated with the same soil-derived bacterial community.

### 2 Media

#### 2.1 Defined 1/2x Taub medium

Previous studies of synthetic CES used a fresh water mimic designed by Taub and Dollar[41] with undefined carbon and nitrogen sources (proteose peptone)[42, 43]. We used the same base medium with chemically defined carbon (glucose) and nitrogen (ammonium) sources in place of the proteose peptone. We modified the medium by adding a stronger phosphate buffer to reduce changes in pH over the course of the experiment. The chemical composition of the medium is shown in Table S4. Media were always prepared no more than two days prior to use. The medium is designed to be carbon limited and the nutrient budget for each CES (including gasses) is given in Table S5.

**Algal growth media:** Prior to the start of an experiment *Chlamydomonas reinhardtii* was grown in Tris-Acetate-Phosphate (TAP) medium following a standard recipe `https://www.chlamycollection.org/methods/media-recipes/tap-and-tris-minimal/`.

### 3 Protocol for initiating experiment

#### 3.1 Algal culturing protocol

A *C. reinhardtii* culture in TAP medium was initiated from a single frozen stock in a 150 mL Erlenmeyer flask containing 10 mL of medium. Cells were grown at 225 rpm shaking and approximately 3000 lux illumination for approximately 5 d. The liquid culture was transferred to a 15 mL sterile Falcon tube and centrifuged at 5000rpm for 2 minutes. The supernatant was quickly discarded, and the pellet was re-suspended in ∼5 mL of 1/2x Taub media described above. The density of algae in the resulting suspension was then measured via hemocytometry. This suspension was then used to initiate the CES where algae were always diluted to a starting density of $5 \times 10^5$ cells/mL.

#### 3.2 Initiating closed ecosystems

All manipulations were performed in a biosafety cabinet. Vials (nominal volume 40 mL CG-4902-08, ChemGlass) and stir bars were sterilized by autoclaving. Each vial was filled with 19.5 mL 1/2x Taub minimal medium. Each vial was then inoculated with 0.5 mL of soil-derived bacterial community and volume of algae yielding $5 \times 10^5$ cells/mL final density (typical volumes $<$0.1 mL).

The metal housing of the pressure sensor, which is mounted on the inside of the vial cap (Figure 1), absorbs light and confounds readings. The manufacturer advises shielding the sensor from direct illumination. To accomplish this, we placed a porous foam stopper ∼1 cm above the meniscus of the liquid. The open-cell foam stopper was cut to size by hand and sterilized by autoclaving. Stoppers shaded the pressure sensor while permitting rapid gas exchange. The foam stoppers also significantly reduced condensation on the sensors. Before the foam stoppers were used, heavy condensation formed in some of the sensors, causing sensor failure. Vials were then fitted with customized metal, plastisol-lined, caps (Burch Bottle and Packaging, `burchbottle.com`, 24-400 black metal plastisol lined cap P/N 3CPLB0241PW) fitted with pressure sensors as described below. Caps were screwed on tightly by hand and wrapped in parafilm. The light intensity was set to 800 Lux (as measured at the top of the aluminium block) in all systems (average error ∼1 %). See section 4.2 for details on light intensity.

##### 3.3 Protocol for CES dilution between rounds of enrichment

Between each round of enrichment each CES was opened and transferred into a 50mL Falcon tube in a biosafety cabinet to ensure sterility. The contents of the Falcon tube were homogenized by pipetting and vortexing. 1 mL of the CES was then transferred to a sterile and clean vial already containing 19mL of 1/2x Taub minimal medium, a sterile foam stopper was inserted into the vial, the cap was again placed on the vial, tightened by hand, wrapped in parafilm and the CES was returned to the same custom culturing device and the experiment was continued. Dilutions occurred either at the end of the light phase or the end of the dark phase.

#### 4 Custom culturing devices

Devices are identical to those presented in a previous study[44] with two modifications: (1) communities were hermetically sealed with plastisol lined metal caps that were retrofitted with pressure sensors that were readout via a RaspberryPi and (2) light intensity from the LED below the vial was attenuated by screens rather than plastic neutral density filters as the latter were found to degrade on the timescale of many months. Below we document these two modifications, including the calibration of light intensity incident on the CES. A schematic of these devices is shown in Figure 1C of the main text. A key feature of these devices is that they permit feedback temperature control of the vial. Each vial fits snugly in a metal block which is under constant feedback control via a Peltier element and thermometer[45]. The feedback temperature control allows for large changes in illumination intensity without changes in pressure due to absorption of light and heating. To demonstrate that these devices alleviate pressure changes driven by heating due to illumination we performed a control experiment with only water in the vial, the result is shown in Figure S1 indicating negligible change in pressure due to light absorption or convective heating the from the LED below the vial.

##### 4.1 Integration of pressure sensors into hermetically sealed vials

Plastisol lined metal caps, compatible with the vials used in our study, were used following the work of Taub and co-workers who reported that plastisol lined metal caps performed the best in terms of hermetic sealing[46]. The pressure sensors used in this study were Bosch BME280 integrated temperature, humidity, pressure sensors on a single small PC board which were purchased from Amazon (ASIN: B0118XCKTG). These small boards fit within the caps on our vials (diameter 1 inch). However, to readout pressure from these sensors requires connecting 4 leads to a RaspberryPi computer. To accomplish this without sacrificing the hermetic seal by the metal caps we developed the following protocol.

A strip of four header pins, which fit the holes in the PC board housing the pressure sensor, were purchased. We then punched a hole in each metal cap with sufficient clearance to allow the header pins to pass through the hole in the cap. The header pins were then fed through the hole in the cap and held in place with modeling clay. We then used a specialized epoxy (EPO-TEK, H74, Epoxy technology) designed for hermetic sealing applications. The epoxy was spread liberally on the outside of the cap as to form a hermetic seal around the header pins while holding them in place. The caps were then placed in an oven at 100 °C for approximately one hour to cure. The caps were then left to finish curing at room temperature for two days, as recommended by the epoxy manufacturers.

The BME280 pressure sensor board was then soldered to the header pins inside the cap. To read the pressure the four leads were connected to the appropriate pins on a RaspberryPi computer to enable I2C communication. We used a Python API developed by Adafruit (`https://www.adafruit.com/`) to acquire data from the BME280 (`https://github.com/adafruit/Adafruit_CircuitPython_BME280`).

The data acquisition was controlled by a custom written Python script which read out the pressure sensor, performed feedback temperature control and controlled the illumination provided by the LED.

##### 4.1.1 Validation of hermetic sealing of vials

To test the quality of the hermetic seal of our caps we performed an experiment where six vials were filled with 20 mL of water, sealed as described above (except the use of foam stoppers), weighed and stored in a incubator room at 30 °C. Vials were then weighed on a precision balance five times over a period of 60 d. We assume any loss of mass to be due to water evaporation. We performed linear regression on the change in mass with time and observed an average loss rate $0.09 \pm 0.14\,\mathrm{mg\,d^{-1}}$. At this rate we expect a CES to lose roughly 4 mg in a 50 d experiment or 0.02 % of its mass. These leakage rates are comparable to those observed in previous CES experiments[43].

#### 4.2 Calibration of light intensity

The LEDs providing illumination were identical to those used in a previously published study from our group[44]. Due to the proximity of the LEDs to the vial and the relatively low intensity used, we needed to attenuate the light. Previous attempts to do this with neutral density filters revealed that such filters slowly degrade over time resulting in changing light intensities on the timescale of months. To solve this problem we instead used metal mesh, placed between the LED and the vial housing the CES (Figure 1, main text). The used metal mesh are 304 Stainless steel wire cloth discs with a hole diameter (D factor) of 0.0021 inches. We placed two layers of this metal mesh between the LED and the vial to achieve the desired range of incident light intensities. The metal mesh was purchased from McMaster-Carr.

To calibrate each of our 8 culturing devices a script was written to slowly vary the LED light intensity by varying a control voltage - from maximum, to zero and back to the maximum level. A lux-meter (Technical Light meter PCE-LED 20 by PCE Americas Inc.) was placed at the top of the metal block (without a vial present) and the measured values were recorded at each set point. Care was taken to allow the LED to equilibrate after each time the light intensity was changed. For each of the 8 systems we fit a polynomial (6th order) to these data to obtain a function $V_{cntl} = f(I)$ where $I$ is the measured intensity and $V_{cntl}$ is the control voltage applied to the LED driver (Buckpuck, 3021, 350mA, `www.ledsupply.com`).

We then quantified the reliability of our calibration by writing a script that used the fits to calculate the control voltage needed to changed the light intensity of each LED to target values. The measured light intensities ($I_{meas}$) were then compared to the target light intensities ($I_{set}$). We then computed an error as $(|I_{set} - I_{meas}|)/I_{set}$ as a function of $I_{meas}$, which we found to be of order 1 % for all systems (Figure S22).

As noted by Mickalide and Kuehn[44] the intensity measured at the top of the metal block is 10-fold lower than the mean intensity experienced by a cell in the vial. Therefore, we expect the mean intensity in the vial (neglecting scattering from cells) to be 8000 lux or approximately $150\,\mathrm{\mu mol\,m^{-2}\,s^{-1}}$. The conversion from lux to $\mu$ mol $\mathrm{m^{-2}s^{-1}}$ was done by measuring the intensity at the top of a metal block in one system using a LI-COR LI-250A light meter with a quantum sensor.

### 5 Pressure data analysis

#### 5.1 Converting changes in pressure to production (consumption) of $CO_2$ ($O_2$)

The air pressure reflects gaseous composition changes in the vial. By ideal gas law,

$$\Delta P = \frac{RT}{V_g}(\Delta n_g(O_2) + \Delta n_g(CO_2)) \tag{S1}$$

where $R$ is the gas constant and $T$ is the CES temperature. Subscripts $g, l, t$ denote quantities associated with *gas*, *liquid* or *total* quantities in the vial (for example, $V_g$ is the gas volume; $n_g(O_2)$ is the number of moles of gaseous $O_2$; and so on). $\Delta n_g(O_2)$ and $\Delta n_g(CO_2)$ are related through photosynthesis/respiration and the individual equilibrium of $O_2$ and $CO_2$ between their respective liquid and gas phases. Our objective is to quantitatively relate the change in pressure to the change in $O_2$ and $CO_2$ in the vial. We begin by noting:

1. In photosynthesis (or respiration), define photosynthetic (respiratory) quotient $\nu$ as the ratio of $O_2$ produced (consumed) and $CO_2$ consumed (produced):

$$\nu = \frac{|\Delta n_t(O_2)|}{|\Delta n_t(CO_2)|} = -\frac{\Delta n_t(O_2)}{\Delta n_t(CO_2)} \tag{S2}$$

   We assume that the rate of $O_2/CO_2$ production and consumption by photosynthesis or respiration is much slower than the equilibration of $O_2/CO_2$ between gas and liquid and the carbon equilibria in water. The fact that our CES are well mixed makes this assumption reasonable. In this limit, the CES always quickly comes to new equilibrium with any $O_2/CO_2$ production or consumption, so the $O_2/CO_2$ produced or consumed is reflected by the total $O_2/CO_2$ change in the CES. Also note that $n_t(CO_2) = n_g(CO_2) + n_l(CO_2)$, where we use $n_l(CO2)$ to denote all forms of dissolved $CO_2$, including $H_2CO_3^*, HCO_3^-$ and $CO_3^{2-}$ molecules ($H_2CO_3^*$ denotes both $CO_{2_{(aq)}}$ and $H_2CO_3$; see section 5.4 for a detailed discussion).

2. The total $O_2$, both gaseous and dissolved, can be calculated by Henry's law:

$$n_l(O_2)/V_l = [O_2]_l = H_{O_2}P_{O_2} = H_{O_2}\frac{RTn_g(O_2)}{V_g} = H_{O_2}RT[O_2]_g \tag{S3}$$

   where $H_{O_2}$ is the Henry's constant and $P_{O_2}$ is the partial pressure of $O_2$. Define

$$u_{O_2} = \frac{\Delta n_l(O_2)/V_l}{\Delta n_g(O_2)/V_g} = H_{O_2}RT \tag{S4}$$

   (the ratio of dissolved $O_2$ concentration and gaseous $O_2$ concentration), and

$$\Delta n_t(O_2) = \left(1 + \frac{V_l}{V_g}u_{O_2}\right)\Delta n_g(O_2). \tag{S5}$$

3. The total $CO_2$ includes gaseous $CO_2$ and dissolved $H_2CO_3^*, HCO_3^-, CO_3^{2-}$ ($H_2CO_3^*$ denotes both $CO_{2(aq)}$ and $H_2CO_3$; see section 5.4 for a detailed discussion):

$$n_t(CO_2) = n_g(CO_2) + n_l(CO_2) = n_g(CO_2) + V_l([H_2CO_3^*] + [HCO_3^-] + [CO_3^{2-}]) \quad \text{(S6)}$$

$$[H_2CO_3^*] = H_{CO_2}P_{CO_2} = H_{CO_2}\frac{RTn_g(CO_2)}{V_g} \quad \text{(S7)}$$

$$H_2CO_3^* \rightleftharpoons H^+ + HCO_3^- : k_a = \frac{[H^+][HCO_3^-]}{[H_2CO_3^*]} \quad \text{(S8)}$$

$$HCO_3^- \rightleftharpoons H^+ + CO_3^{-2} : k_2 = \frac{[H^+][CO_3^{2-}]}{[HCO_3^-]} \quad \text{(S9)}$$

Similarly, define $u_{CO_2} = \frac{\Delta n_l(CO_2)/V_l}{\Delta n_g(CO_2)/V_g}$, and

$$\Delta n_t(CO_2) = \left(1 + \frac{V_l}{V_g}u_{CO_2}\right)\Delta n_g(CO_2), \quad u_{CO_2} = H_{CO_2}RT\left(1 + \frac{k_a}{[H^+]} + \frac{k_a k_2}{[H^+]^2}\right) \quad \text{(S10)}$$

Using the formalism developed above we can compute the *total* change in $CO_2$ from a measured change in pressure as follows:

$$\Delta P = -\frac{RT}{V_g}\left(\frac{\nu}{1 + (V_l/V_g)u_{O_2}} - \frac{1}{1 + (V_l/V_g)u_{CO_2}}\right)\Delta n_t(CO_2), \quad \text{(S11)}$$

$$\text{where} \quad u_{O_2} = H_{O_2}RT, u_{CO_2} = H_{CO_2}RT\left(1 + \frac{k_a}{[H^+]} + \frac{k_a k_2}{[H^+]^2}\right)$$

We refer to $\frac{\Delta P}{\Delta n_t(CO_2)}$ as the conversion factor of $n_t(CO_2)$. Note the sign which indicates that a decline in pressure results from the production of $CO_2$ and consumption of $O_2$. Further recognize that $u_{CO_2}$ depends on the $pH$ of the water through the impact of the $pH$ on the $CO_2$ equilibria.

When calculating conversion factors, we also account for chemical constants' dependence on temperature.

$$\text{Henry's constant [47]:} \ \ln(H) = A + B/T + C\ln(T) \quad \text{(S12)}$$

$$\text{Equilibrium constants [48]:} \ pK_T = pK_\theta + \frac{1}{R\ln 10}\left(\Delta_r H_\theta^\circ\left(\frac{1}{\theta} - \frac{1}{T}\right) + \Delta_r C_{p\theta}^\circ\left(\frac{\theta}{T} - 1 + \ln\frac{T}{\theta}\right)\right) \quad \text{(S13)}$$

where $A, B, C$ are parameters for Henry's constants, $pK = -\log_{10} k$, $\Delta_r H^\circ$ is the standard enthalpy of reaction, $\Delta_r C_p^\circ$ is the standard heat capacity of reaction, and $\theta = 298.15K$.

Table S3 summarizes parameters and chemical constants used in the calculation. All constants on the RHS of Equation S11 are known or have been measured with the exception of $\nu$ which we assume to take a value of 1.

#### 5.2 Comparing with $O_2$ measurement

To validate the pressure measurement we performed a control experiment with a CES where we measured pressure and $O_2$ levels concurrently. The conversion factor between pressure and $O_2$ concentration can be found by combining Equation S2 and S5 and substituting in Equation S11,

$$\Delta P = RT\left(1 - \frac{(1+(V_l/V_g)u_{O_2})}{\nu\,(1+(V_l/V_g)u_{CO_2})}\right)\Delta[O_2]_g, \tag{S14}$$

$$\text{where}\quad u_{O_2} = H_{O_2}RT, u_{CO_2} = H_{CO_2}RT\left(1+\frac{k_a}{[H^+]}+\frac{k_a k_2}{[H^+]^2}\right) \tag{S15}$$

We refer to $\frac{\Delta P}{\Delta[O_2]_g}$ as the $O_2$ conversion factor. $O_2$ levels (concentrations) were measured non-invasively using a Presens (`https://www.presens.de/`) luminescence quenching based method. We used a PSt3-YAU autoclavable sensor spot which was adhered to the inside of one of our vials using optical glue as per the manufacturer instructions. We integrated the optical fiber into one of our custom culture devices (Figure 1, main text). We then made short term measurements of both pressure and oxygen and the results are shown in Figure S2, panel A-B.

Figure S2 C-D shows pressure verses $[O_2]$ where we observe the expected linear dependence. However, the measured slope differs from theoretical prediction at $pH = 6.5, \nu = 1$ given by equation S14. The difference between the measured slope and our theoretical prediction can be accounted for by changes in $pH$ and $\nu$.

#### 5.3 Corrections to conversion factors

The $O_2$ conversion factor (Equation S14) explicitly depends on three quantities: temperature $T$, pH and photosynthetic/respiratory quotient $\nu$ (Figure S3). The dependence on temperature is weak, and with the system under temperature control at 30 °C, temperature fluctuations are small (∼0.1 °C, Figure S2 A-B). The dependencies on pH and $\nu$, however, are strong.

The measured $O_2$ conversion factor differs from the theoretical prediction at $pH = 6.5, \nu = 1$. This can be explained by the fact that we cannot continuously measure $pH$ and $\nu$. Figure S3 B shows that the measured values correspond to a region in the $(pH, \nu)$ space that is reasonable for the CES organisms and environmental conditions [49]. The measured conversion factor also changes between cycles and between light/dark conditions (figure S2 D). This can arise from dynamical changes of $pH$ and $\nu$ due to different metabolic activities at different time and light/dark conditions. For example, the photosynthetic and respiratory quotients are likely not identical. Drift in the $O_2$ measurement by the Presens sensor could also give rise to these changes.

It is conceivable that the conversion factors can be further corrected by other contributions. However, we consider these contributions to be either negligible compared to effects of $pH$ and $\nu$, or not quantifiable given our knowledge of the system.

- Water vapor pressure was neglected in Equation S1 because its contribution is negligible. The partial pressure of water vapor is

$$P_{H_2O} = P_{sat} * RH \tag{S16}$$

where $P_{sat} = 4.247KPa$ (the saturation pressure at 30 °C[50]) and $RH$ is the relative humidity. We measure RH using the BME280 pressure sensor and observe that RH rises in the first few hours of the experiment and then remains stable with small fluctuations (Figure S2 A-B). We also examined RH during light-dark cycles for each CES during each round of dilution. Approximately 80 % of the CES show no measurable change in RH due to LED illumination. In those CES where appreciable change in RH occurred, fluctuations were <0.4 % which correspond to change in pressure of approximately 0.2hPa. Given that changes in pressure due to $O_2$ levels are typically between 4hPa and 10hPa we conclude that the contribution of water vapor to our carbon cycling measurements is negligible.

- Ions (such as $Ca^{2+}$ and $Mg^{2+}$) in the solution affect Henry's law constants [47] and carbonate equilibrium. However, because ions are constantly utilized by organisms, we cannot quantitatively model their effects.

#### 5.4 More details about carbonate equilibria in water

$CO_2$ dissolves in water by three steps:

$$CO_{2(aq)} + H_2O \rightleftharpoons H_2CO_3 : \; k_{a1} = \frac{[H_2CO_3]}{[CO_2]_{(aq)}} \tag{S17}$$

$$H_2CO_3 \rightleftharpoons H^+ + HCO_3^- : k_{a2} = \frac{[H^+][HCO_3^-]}{[H_2CO_3]} \tag{S18}$$

$$HCO_3^- \rightleftharpoons H^+ + CO_3^{-2} : k_2 = \frac{[H^+][CO_3^{2-}]}{[HCO_3^-]} \tag{S19}$$

We adopt the convention of using $H_2CO_3{}^*$ to denote both $CO_{2_{(aq)}}$ and $H_2CO_3$ and using an apparent equilibrium constant to combine S17 and S18:

$$H_2CO_3{}^* \rightleftharpoons H^+ + HCO_3^- : k_a = \frac{[H^+][HCO_3^-]}{[H_2CO_3^*]} = \frac{[H^+][HCO_3^-]}{[H_2CO_3 + CO_2]} \tag{S20}$$

See [51] for a detailed discussion. This is the same convention in [50] (adopted from [52] and [48]), where we adopted all equilibrium-related numbers. The Henry's constant is only slightly affected by this convention:

$$H^*(CO_2) = \frac{[H_2CO_3^*]}{P_{CO_2}} \approx H(CO_2) = \frac{[CO_{2_{(aq)}}]}{P_{CO_2}} \tag{S21}$$

because $[CO_{2_{(aq)}}] \gg [H_2CO_3]$.

#### 5.5 Calculating carbon cycling rate

To compute the number of moles of carbon cycled per day we first compute the rate of respiration $r$ during the dark phase by linear regression (Figure S6) and the application of Equation S11 using $pH = 6.5$ (measured pH at the end of all rounds of enrichment) and $\nu = 1$. We assume the rate of respiration $r$ is constant during light and dark phases and compute the total number of moles $CO_2$ respired in a light-dark cycle as $n_r^{tot} = r \times 24h$. We then compute the *net* number of moles of $CO_2$ fixed during the light phase ($f$, Figure 1, main text) by measuring the change in pressure over the course of the light phase and again applying Equation S11, yielding $n_f \propto \Delta P_{light}$ - the change in pressure during the light phase. To compute the total number of moles $CO_2$ fixed during the light phase we account for the respiration that occurred during the light phase by adding $n_f^{tot} = n_f + r \times 12h$. The result is a quantification of the total number of moles of $CO_2$ fixed ($n_f^{tot}$) and respired ($n_r^{tot}$). We then compute the number of moles cycled per day as:

$$n_c = min(n_f^{tot}, n_r^{tot}) \tag{S22}$$

We compute $n_c$ for each light-dark cycle and the results are shown in Figure 2 of the main text.

### 6 Further analysis of pressure measurements

#### 6.1 Minimum detectable change in pressure

We estimated the minimum detectable change in pressure in a 12 hour period by examining the dark phase pressure dynamics across all dark phases during all four rounds of enrichment. For each dark phase we fit a 3rd order polynomial least-squares fit to the pressure decline. The residuals to this fit contained no observable temporal dynamics by eye and had zero mean on average. As a result, these residuals quantify the noise in the pressure measurement itself. The standard deviation of these residuals ($\sigma_p$) agreed well with the short timescale (1 h) fluctuations in pressure in the water-only control experiment (Figure S1). The median $\sigma_p$ across all dark phases, systems and CES was 0.095hPa (5th and 95th precentiles: 0.055hPa and 0.26hPa, respectively). These fluctuations set the minimum detectable change in pressure. To approximate this minimum detectable change in pressure we estimated the uncertainty in the pressure given pressure fluctuations of order 0.095hPa and a measurement time of 12 h (the duration of one light or dark phase). We first computed the autocorrelation time of presssure fluctuations to be $\sim$3 min on average across all dark phases, rounds and CES. Therefore, in a given 12 h period there are 240 statistically independent measurements of the pressure. Thus, the minimum detectable change in pressure is of order $\Delta p_{min} = 0.095/\sqrt{240} = 0.0061$ hPa. Above we compute the number of moles of $CO_2$ fixed or produced per unit change in pressure to be: $1.2821 \times 10^{-6}$moles/hPa, which yields a minimum detectable change in $CO_2$ of approximately $7.8 \times 10^{-9}$ moles.

To understand the magnitude of this number we compute the number of *E. coli* cells that can be produced given $7.8 \times 10^{-9}$ moles of C atoms available for biomass. The number of C atoms per cell of *E. coli* is approximately $7 \times 10^9$ (`https://bionumbers.hms.harvard.edu/bionumber.aspx?&id=103010`. $7.8 \times 10^{-9}$ moles of C yields approximately $6.7 \times 10^5$ *E. coli* cells. In our culture volume of 20 mL this corresponds to a density of only $3.3 \times 10^4$ cells/mL which a very low density for bacteria in culture!

#### 6.2 Respiration rates during the dark phase

During the dark phase we compute the respiration rates by first converting our pressure measurement to an increase in $CO_2$ levels within the CES using the formalism derived above and then fitting a line to the decline in pressure that occurs during the dark phase. We find that a constant respiration rate during the dark phase (e.g. purely linear decline in the pressure during the dark phase) is a good approximation to our data. To quantify this we fit linear and quadratic polynomials to the decline in pressure we observe during the dark phases (Figure S6, top row). We then compute the residual for both linear and quadratic models and compute the ratio of the residuals $\sigma_{res}^{linear}/\sigma_{res}^{quad}$. In the case where the decline in pressure is purely linear, and therefore the respiration rate constant throughout the dark phase, we expect the linear and quadratic fits to the data to be nearly identical and hence $\sigma_{res}^{linear}/\sigma_{res}^{quad} \approx 1$. Figure S6 (bottom four panels) show $\sigma_{res}^{linear}/\sigma_{res}^{quad}$ as a function of time for all four rounds of dilution. We find that the linear model is a good one for describing the decline in the dark phase pressure for nearly all of the data. Note that even for $\sigma_{res}^{linear}/\sigma_{res}^{quad} \approx 2$ the departure from linearity is small Figure S6 (top left panel).

#### 6.3 Transient decline in pressure during round 1

In Figure 2B of the main text we show a time series of pressure during round 1 for a single CES. Identical traces for all CES in round 1 are shown in Figure S5. We note that for 7 of 8 CES we observe a relatively abrupt drop in pressure around 25 days after closure. The exception being CES B.3 (Figure S5).

To understand the reason for this decline we analyzed the pressure data in more detail. First, we estimated the rate of $O_2$ production during the light phase of each light-dark cycle. To accomplish this we performed a spline regression on the pressure as a function of time during each light phase. We used the 'fit' function in MATLAB which optimizes an objective function: $w\sum(p_i - s(t_i))^2 + (1-w)\int(\frac{d^2s(t)}{dt^2})^2dx$ where $s$ is the piece-wise cubic fit to the data and the integral in the second term is evaluated over the domain of the data. The $p_i$ and $t_i$ correspond to our data. We used $w = 0.8$ for all fits which avoided fitting short timescale (minutes) pressure fluctuations. An example of an smoothing spline applied to our data is shown in Figure S7A. From these smoothing splines we can directly estimate $\frac{dp}{dt} \propto \frac{dO_2}{dt}$ (see derivation above for this conversion) and an example is shown in Figure S7B. Using this approach we estimated the net $O_2$ production rate by the algae during the light phase. In this calculation we neglected the $O_2$ consumption due to respiration during the light phase. We next plotted $\frac{dO_2}{dt}$ during the light phase for each light-dark cycle that occurred during round 1 (Figure S7D). We find that concomitant with a decline in overall pressure we observe a slowing oxygen evolution rate by the algae during the light phase (compare Figure S7D day 20 to day 30). We also computed the respiration rate during each dark phase via linear regression (e.g. Figure S6A,B) and the results are shown in Figure S7E. We observe that the decline in $\frac{dO_2}{dt}$ during the light phase is accompanied by an increase in the respiration rates during the corresponding dark phases. These two observations suggest that the drop in pressure could arise from a loss of algal biomass (e.g. via senescence) which produces organic carbon that is consumed by the bacterial community. The eventual stabilization of the pressure at longer times suggests a homeostatic mechanism may stabilize the CES e.g. by simulating algal recovery due to higher $CO_2$ levels. A detailed investigation of this phenomenon is beyond the scope of our study.

##### 6.4 Comparison of $O_2$ production rates to literature values

As a means of externally validating our measurement of $O_2/CO_2$ production consumption we make comparisons to available data on *C. reinhardtii* in the literature. Vejrazka *et al.* measure the rate of $O_2$ production per gram biomass (Figure 4 of Ref[53]). There they find that at an intensity of 200 $\mu$mole $m^{-2}s^{-1}$ (approximately our light level) the net oxygen production rate is 1 $\mu$mole $s^{-1}g^{-1}$.

We can estimate an upper limit on the oxygen production rate by the algae. Assume that all of the available carbon is locked up in algal biomass. Assuming a carbon fraction of biomass (dry weight) of 0.5 [54] implies that we have at most approximately $5 \times 10^{-3}$ g dry weight in algal biomass. At the estimated illumination in our system this would correspond to oxygen production rates of about 18 $\mu$mole $h^{-1}$. For comparison, with our data we observe *net* $O_2$ production rates peak at about 4 $\mu$mole $h^{-1}$ (Figure S7D). If we assume dark phase respiration rates (Figure S7E) are sustained during the light phase, we expect that total $O_2$ production by algae is around 5 $\mu$mole $h^{-1}$. We note that this number is well below the maximum rate estimated from the literature and biomass estimates above. This difference arises due to the fact that not all C atoms are in algal biomass. Overall, this estimate provides additional confidence in our pressure based metabolic measurements.

### 7 Metabolic assays

#### 7.1 Ecoplate carbon source respiration assay

Ecoplates were purchased from Biolog (www.biolog.com) and used without modification. After each round of dilution the contents of each CES were homogenized by rapid vortexing and pipetting up and down using a serological pipette. 1.5 mL of homogenized CES was then mixed with 13.5 mL of a modified version of 1/2x Taub minimal media. The media used for this assay lacked any carbon but still had all other compounds present in the same proportions as the complete media (see Table S4). The 1-to-10 diluted CES were then aliquoted into the Ecoplates (100 µL per well). Plates were then wrapped in parafilm and incubated at 30 °C, 250 rpm shaking. To avoid evaporation over the course of the experiment each plate was sealed in a ziplock plastic bag with a moist paper towel. Optical density measurements for each plate were made daily for four days using a BMG Labtech Clariostar plate reader. The $OD_{590}$ values were used without background subtraction and examples are shown in Figure 3 of the main text.

##### 7.1.1 Analysis of ecoplate data

Ecoplate data consisted of time series of dye absorbance like the ones shown in Figure 3C of the main text. Each time series was analyzed as follows: Let $Abs_{i,r}(s)$ denote the time series of $OD_{600}$ measurements where $s$ is the sampling time in hours $s \in \{0, 24, 48, 72, 96\}$.

- (1) Compute $\Delta A_1 = abs(max(Abs_{i,r}) - Abs_{i,r}(0h))$
- (2) Compute $\Delta A_2 = abs(Abs_{i,r}(96h) - Abs_{i,r}(0h))$
- (2) if $\Delta A_1 < h_1$ or $\Delta A_2 < h_1$ then $\tau \to \infty$.
- (3) otherwise, linearly interpolate the data and compute:
  (3.1) $\tau_{i,r}$ such that $Abs_{i,r}(\tau_{i,r}) = h_2 \times max(Abs_{i,r})$

In all of the analysis presented here we chose $h_1 = 0.3$ and $h_2 = 0.9$. The analysis results in a vector of $\tau$ for respiration of each carbon source in triplicate. We then use the quantity $1/\tau$ as a measure of the rate of carbon utilization.

Each CES was thus associated with three vectors, corresponding to the three replicate measurements, of length 32 of $1/\tau$ values for the 32 carbon sources. Each element of these vectors is the consumption rate of a particular carbon source. Let $T^d_{c,r}$ represent such a vector length 32 for the $r^{th}$ replicate of the $c^{th}$ CES at dilution round $d$. Here $r \in \{1, 2, 3\}$, representing the three replicates, and $c \in \{1, 2, 3, ..., 8\}$, representing the eight CES, and $d \in \{1, 2, 3, 4\}$, representing the four dilution rounds. The standard deviation in the consumption rates of each carbon source was computed across all eight CES and all three replicates (24 measurements in total) at each dilution round, resulting in a vector of length 32 for each dilution round whose elements were the standard deviation in the rate of consumption for a particular carbon source at that dilution round. This vector, $V$, is given by Equation S23, where the superscripts of $\sigma$ indicate the quantities over which the standard deviation is computed.

$$V^d = \sigma^{i,r}(T^d_{i,r}) \tag{S23}$$

The distribution of these 32 standard deviations for each dilution round, is plotted in Figure S20. The higher these standard deviations, the greater the differences in $1/\tau$ across CES. The lower the standard deviation, the more similar the rates of carbon utilization across CES. The tests for significance were done by bootstrapping as described in section 7.1.2.

##### 7.1.2 Bootstrapping

To compare the distributions of standard deviations ($V^d$) and JSD (see section 9), bootstrapping was used. In particular, the two distributions being compared were re-sampled with replacement 10,000 times. The medians were computed for each re-sampled distribution, and the difference in medians for the two distributions were found. The p-values were the fraction of the differences that were negative.

##### 7.1.3 Gas Chromatography - Mass Spectrometry (GC-MS)

To quantify those carbon compounds that the algae can excrete we performed gas chromotography-mass spectrometry on algal spent media. These experiments were performed in the absence of any bacteria and in open vessels and these important distinctions may alter algal excretions relative to the CES. However, the experiment does demonstrate some carbon compounds that the algae can excrete.

The lab strain *C. reinhardtii* was grown from frozen stock in TAP medium, with constant shaking and illumination. After 3 days of growth, the cells were washed (centrifuging at 1000rpm for 2 min) in the experimental medium (1/2X Taub prepared as described before, with additional 3.1 mM phosphate buffer, 8 mM $NH_4Cl$ and 8 mM Carbon from glucose). The cells were diluted and re-suspended in the experimental medium in 3 autoclaved vials with sterile stir bars in them, so that the cell density was $10^6$ cells/mL. The vials were capped off with sterile foam stoppers that allow gas exchange with the atmosphere. The vials were placed in metallic casings which were temperature controlled via Peltiers as described above. The metallic casings were illuminated from below with LEDs with the same spectrum as those used the CES experiment at an intensity of approximately 10 000 lux ($\sim 187 \mu$mol $^{-2}$ s$^{-1}$), during the light phase of 12 h-12 h light-dark cycles. After three days of growth, $500 \mu L$ samples were collected from all three vials and centrifuged at 7000 rpm for up to 15 min to ensure all cells were pelleted. The supernatant was collected and stored at $-20\,^{\circ}$C. This procedure was repeated after 6 days, 9 day and 12 days. The collected samples and a sample of the medium were sent to the Roy J Carver Biotechnology Center at UIUC for GC-MS analysis (Agilent 7890A GC/5975C MS). The results, with the GC-MS signal from fresh medium subtracted, are in Supplementary Data 3.

To find the compounds which are excreted in significant amounts, we did a linear regression using least squares on the GC-MS peak height verses day of spent media extraction. The results of the regression provide the slope and p-value, for the null hypothesis that the slope is zero. We defined a compound to be excreted in significant amounts if the p-value was below 0.05, and if the slope was positive, and if all the data were positive. The last condition was necessary because some compounds were present in the medium, but not in the samples resulting in negative data. The significantly excreted compounds are listed in table S6. The second column of the table lists the compounds in the Ecoplate used to measure carbon utilization profiles that are similar to the compounds excreted by the algae.

#### 7.2 Microresp assay for determining nutrient limiting respiration

After each round of enrichment we performed an assay to determine the nutrient limiting respiration in each CES. To accomplish this we used the microresp™(`https://www.microresp.com`). Briefly, microresp measures the production of $CO_2$ during respiration in the dark. The platform uses two 96-well plates, one deep-well (well volume 1.2 mL) and one standard “indicator” plate. The sample is placed in the deep-well plate which is sealed (face-to-face using a custom rubber gasket) with the indicator plate. The indicator plate contains a pH sensitive dye in an agarose gel. Consumption of any available organic carbon via respiration in each well produces $CO_2$ which reduces the pH in the indicator gel as it is absorbed. The $CO_2$ production can then be assayed by removing the indicator plate and rapidly performing absorption measurements on a plate reader.

**Details of microresp assay and calibration:** Each well of the indicator plate is filled with 150 µL of the indicator gel which contains 34.9 µM cresol red, 168.7 mM KCl, 2.81 mM sodium bicarbonate and 3 % agarose. The indicator solution is loaded into the plates at ∼60 °C and then allowed to cool at room temperature for ∼20 minutes. Plates are stored in a sealed ziploc bag with a beaker of water to prevent drying of indicator gels and a beaker of soda lime to prevent $CO_2$ contamination.

To calibrate the $CO_2$ production we performed a control experiment using *Escherichia coli* in a carbon-limited M9 minimal medium with varying levels of glucose from 1.25 mM to 10 mM. Cells were allowed to grow for 24 h and an absorbance spectrum of the gel was measured using a BMG Clariostar plate reader. From these data we determined that the ratio of absorbances at two wavelengths scaled like a power law with the available glucose. Namely,

$$\frac{Abs_{430nm}}{Abs_{570nm}} \propto [Glu]^{1/5} \tag{S24}$$

Under the assumption that the glucose is converted to $CO_2$ with fixed fraction (carbon use efficiency) by the *E. coli* under the range of conditions tested (we expect this to be true and no fermentation to occur) then we can assume that $[CO_2] = \gamma[Glu]$ where $\gamma$ has not been measured here. Under this assumption $Abs_{430nm}/Abs_{570nm} \propto [CO_2]^{0.2}$ where the contant of proportionality is not known.

We next define the ratio $r_t = Abs_{430nm}/Abs_{570nm}$ for a measurement that occurs at time $t$. In the experiment we take two measurements $r_{0h}$ and $r_{24h}$ and then compute the fractional change in $CO_2$ as follows:

$$F_{CO_2} = \frac{CO_2(t = 24h) - CO_2(t = 0h)}{CO_2(t = 0h)} = \left(\frac{r_{24h}}{r_{0h}}\right)^5 - 1 \tag{S25}$$

Note the equality holds because the unknown constant of proportionality cancels out. Therefore, to determine the fractional change in $CO_2$ in each well of the 96-well plate we measure absorbance prior to sealing the wells and after 24 h and compute the quantity above. The results are shown in Figure S21.

**Assay procedure:** To perform an experiment, CES were opened and 240 µL samples were loaded into 12 wells of a 96-deep well plate. To assay nutrient limitation, these CES samples were amended with an additional 10 µL of media that contained: water, phosphorous, carbon, or nitrogen (each in triplicate). Nutrients were added such that the final concentrations were 10 mM, 8 mM or 4 mM for C, N and P respectively. Three additional wells were loaded with 250 µL water. Dispensing of nutrient additions into the 96-well plate was accomplished with a Formulatrix Mantis liquid handling robot. Wells were arrayed in a checkerboard pattern and no wells on the periphery of the plate were used. This layout was necessary to avoid leakage effects between the wells and with the atmosphere.

An indicator plate was then removed from storage, absorbance was assayed and the plate was clamped tightly to the deep-well plate using the custom rubber gasket and metal clamp. The clamped assembly was incubated in the dark for 24 h with shaking at 250rpm and maintained at 30 °C. After the 24 h, the clamp was removed and the indicator plate rapidly assayed again for absorbance at the two wavelengths. The resulting data was analyzed as described above. Measurements that extended beyond 24 h suffered from substantial well to well leakage confounding measurements.

###### 7.2.1 Results of microresp assay

Microresp assays were performed on all CES after each round. Comparing the fractional change in $CO_2$ produced ($F_{CO_2}$) across samples from the same CES and the same round amended with different nutrients allows us to determine which nutrient is limiting respiration. For example, see the upper left hand panel of Figure S21. Each column shows a different amendment with 'water' indicating no added nutrients. The data in the 'cntl' column are from wells containing only water and indicates the spurious $CO_2$ signal due to leakage and systematic errors of the measurement. Therefore, by examining a single panel, the column with the largest increase in $CO_2$ produced (relative to no added nutrients) is the nutrient limiting respiration. So for example, the microresp assay after round 1 in CES A.2 indicates that respiration is P-limited. In contrast, the microresp assay after round 1 in CES B.1 indicated that respiration is C-limited (Figure S21).

Figure S21 shows all of the microresp data for all CES after all rounds of enrichment. In general, we find that CES from soil sample A exhibit P-limited respiration while those from soil sample B exhibit C-limited respiration. However, the nutrient limiting respiration varies between rounds for CES A.2 going from P-limited at round 1 to C-limited in rounds 2 and 3 and then back to P-limited in round 4. Respiration is never N-limited in our conditions.

To test whether the identity of the limiting nutrient impacted the carbon cycling rate we compared the average carbon cycling rates for CES derived from soil samples A and B. We found no significant difference between average cycling rates (measured on the last day of each round) for CES from samples A and B p-values: 0.53, 0.23, 0.85, 0.67 for rounds 1 to 4 respectively. We conclude that carbon cycling rates are robust to C- and P-limited respiration.

###### 7.2.2 Stoichiometry and P-limitation

We observe P-limitation in CES originating from one of the two soil samples (A). Here we ask whether this P-limitation could have arisen from P incorporated into biomass or not. Typical ratios of carbon to phosphorous in biomass are of order 100:1 (e.g. 100 C atoms for each P atom). Let us assume for a moment that at steady state the vast majority of the available carbon in the system is in biomass. This means that there are $2 \times 10^{-4}$ moles of C in biomass, and roughly $2 \times 10^{-6}$ moles P in biomass. As Table S5 shows there are $8 \times 10^{-5}$ moles of P available in the system at the outset. This suggests that P is in excess by approximately a factor of 40.

This is a rough estimate, so here we solidify it further by looking into the biomass stoichiometry of *C. reinhardtii* and typical bacteria. For *C. reinhardtii* Boyle *et al.*[54] measure C:N ratios are between 5:1 and 14:1 depending on whether the cells are growing autotrophically or heterotrophically (see Table 3, Ref [54]). In a separate study of growth at low temperature, the authors report an N:P ratio of between 26.5 and 36.5[55]. This gives a range of C:P ratios for the algae of 511:1 to 132:1. At these ratios, the upper bound of P held in algal biomass, assuming all of the C atoms are in algal biomass, would be approximately $2 \times 10^{-4}/132 = 1.5 \times 10^{-6}$ moles. This is about a factor 80 below the available P.

For bacteria, typical C:N ratios are vary between 5:1 and 10:1 and C:P ratios around 60:1 to 100:1[56]. Assuming all available C atoms are in bacterial biomass, stoichiometry puts an upper bound of $2 \times 10^{-4}/60 = 3.3 \times 10^{-6}$moles of P. This estimate of moles of P in bacterial biomass is still below the $8 \times 10^{-5}$moles which are available.

These analyses strongly suggest that sequestration of phosphate by either *C. reinhardtii* or bacteria in our CES is responsible for the P-limitation we observe in CES which arise from soil sample A. The molecular basis of this sequestration remains for future work.

##### 7.3 Measurement of pH at the end of the experiment

At the end of each round of the experiment the pH was measured in each CES using litmus paper. For all rounds and all CES the pH was found to be 6.5.

##### 7.4 Measurement of total organic carbon

After each round of enrichment we measured total organic carbon in each CES. To accomplish this, we prepared diluted samples of each CES in a solution of 0.5% v/v phosphoric acid. All samples were sent to the Illinois State Water Survey where they performed measurements of non-purgable organic carbon (NPOC). The survey lab employed a high temperature combustion method (5310B, `https://www.nemi.gov/methods/method_summary/5717/`, lab webpage `https://www.isws.illinois.edu/chemistry-and-technology/analytical-services-laboratory`). Each sample was processed in 5 replicates and outliers were discarded. The mean and standard deviation were computed from at least 3 replicate measurements and are shown in Figure S8. The gray line in Figure S8 represents the organic carbon initially supplied as glucose. There are three other sources of carbon in the CES: initial inoculum of algae, $CO_2$ and biomass in the initial soil sample. The first two of these contributions are negligible and the third has not been quantified.

#### 8 16S Sequencing

##### 8.1 DNA extraction

Qiagen's DNeasy Blood and Tissue Kit (Cat No./ID: 69581) was used to extract DNA from the communities. Pre-treatment with a Lysis buffer was performed to obtain DNA from any gram positive bacteria in the community. The Lysis buffer contains 2 mM Na EDTA made in 20 mM Tris-Cl at pH 8, 1.2% Triton X-100 (v/v) and 20 mg/mL Lysozyme added immediately before use. Lysozyme from chicken egg white (SKU L6876 from SigmaAldrich) was used. Frozen samples were thawed and 250 $\mu$L of the sample transferred to Eppendorf tubes. These were centrifuged at 14000 rpm for 15 min, and the supernatant discarded. The cell pellet was re-suspended in 180 $\mu$L of the previously prepared lysis buffer and incubated in a water bath at 37 C for 30 min. 25 $\mu$L proteinase K and and 150 $\mu$L buffer AL (without ethanol) were added to each tube. The tubes were incubated in a water bath at 56 C for one hour. 200 $\mu$L of ethanol was then added to each tube before vortexing thoroughly. The samples were then transferred to the DNeasy 96 well plate, placed on an S block (provided with the kit), which was then sealed. The plate was then centrifuged in plate centrifuge at 4000 rpm for 15 min. The flow through was discarded, and 500 uL of buffer AW1 (pre-mixed with ethanol) was added to all the wells. The plate was re-sealed and centrifuged at 4000 rpm for 10 min. The flow through was discarded and 500 $\mu$L of buffer AW2 (pre-mixed with ethanol) was added to all wells and centrifuged (without sealing) for 20 min at 4000 rpm. Next, the DNeasy plate was placed on a rack of elution tubes. 100 $\mu$L of elution buffer was added to all wells, the plate was sealed and centrifuged at 4000 rpm for 3 min. The last step was repeated, so as to get 200 $\mu$L of DNA in the elution tubes. The elution tubes were closed with caps and stored at $-20\,^{\circ}$C.

To extract DNA from the initial soil samples, Qiagen's DNeasy Power Soil Pro Kit (Cat No./ID: 47014) was used. Samples were collected after the 48 h growth phase in the dark. The beads of PowerBead Pro tubes were carefully removed and 500 $\mu$L of the soil sample was added to them. They were centrifuged for 30 s at 10000 g. The supernatant was removed and the beads were added back into the tubes. 800 $\mu$L of solution CD1 was added to the tubes and vortexed briefly to mix. The tubes were then horizontally secured to a Vortex Adapter and vortexed at maximum speed for 10 min. The tubes were then centrifuged at 15000 g for 1 min. The supernatant was transferred to a 2 mL microcentrifuge tube. 200 $\mu$L of solution CD2 was added to the microcentrifuge tube and vortexed for 5 s. The mixture was centrifuged at 15000 g for 1 min. The supernatant was transferred to another microcentrifuge tube. 600 $\mu$L of solution CD3 was added and vortexed for 5 s. The lysate was loaded to Spin Columns and centrifuged for 1 min. Once all the lysate passed through the spin columns, they were placed in collection tubes and washed with 500 $\mu$L solution EA by centrifuging for 1 min at 15000 g. The flow through was discarded and 500 $\mu$L solution C5 was added to the Spin Column and centrifuged at 15000 g for 1 min. The flow through was discarded and the Spin Columns placed in fresh collection tubes. The tubes were centrifuged at 16000 g for 2 min and the Spin Columns were placed in elution tubes. 75 $\mu$L of solution C6 was added to the center of the filter membrane and the tubes were centrifuged at 15000 g for 1 min to obtain the DNA in the flow through.

#### 8.2 Library Preparation

##### 8.2.1 DNA quantification

After extraction, DNA was quantified using Qubit dsDNA BR Assay Kit (Catalog number: Q32853 from Thermo Fisher Scientific). Due to the large number of samples, a modified procedure using a plate reader was followed according to [57]. An 8 point standard curve was made by serially diluting the 100 ng/$\mu$L standard with the 0 ng/ $\mu$L standard in one column of a 96 well plate. In another column of the plate, 195 $\mu$L of the Qubit working solution was added to eight wells. 5 $\mu$L of the serially diluted standards are added to the wells containing the working solution. The plate was briefly vortexed to ensure complete mixing of sample and working solution. The plate was then placed in the plate reader. The excitation wavelength was 485 nm and the emission wavelength was 530 nm. Fluorescence was measured for all 8 wells. The fluorescence values were background subtracted, and plotted against the known concentrations of the standards on a log-log scale. A straight line was fit to the data, which resulted in a power law for the standard curve.

A similar procedure was followed to estimate the DNA concentration in the samples. 195 $\mu$L of the working solution was added to all the wells of a 96 well plate. 5 $\mu$L of the samples were then added to the wells. The plate was vortexed briefly and fluorescence measurements taken in a plate reader. Using the previously obtained standard curves, the readings were converted to concentration of DNA.

##### 8.2.2 PCR

The primers created by the Earth Microbiome Project were used for performing PCR [58]. The primers (515F - 806R) target the V4 region of the 16S subunit of the rRNA. The V4 region is approximately 254 bp long. The primers include barcodes, linkers, pads, and adapters. Taking these into consideration, PCR products of about 390 bp were expected. The reverse primers contain unique barcodes that allowed de-multiplexing of reads into communities. All samples received the same forward primer and different and unique reverse primers. Platinum Hot Start

PCR Master Mix (2x) from ThermoFisher (cat. no. 13000014) was used. The reagents were added in the order and volume presented in Table S1 to 96 well PCR plates. Every reaction was performed in triplicate.

The thermocycler settings are in Table S2. The triplicate PCR products were pooled to get a total volume of 75 $\mu$L.The DNA content in the pooled PCR products were quantified using the Quibit assay described above. Once the concentration of the PCR products was obtained, the volume containing 240 ng of DNA from each sample was calculated. This volume was then pooled into a fresh Eppendorf tube. This ensured the amount of DNA to be used for sequencing was normalized.

Qiagen's QIAquick PCR purification (catalog number 28104) was used to clean the pooled PCR products. 5 volumes of buffer PB with a pH indicator was added to 1 volume of the pooled PCR products in QIAquick spin columns placed in collection tubes. The columns were centrifuged at 17900 g for 45 s. The flow through was discarded and 0.75 mL of buffer PE was added. The columns were centrifuged at 17900 g for 45 s. The flow through was discarded and the columns were centrifuged at 17900 g for 1 min to completely remove any residual ethanol. The columns were then placed in clean microcentrifuge tubes. 50 $\mu$L of buffer EB was added to column and was allowed to stand for a minute and then centrifuged at 17900 g for 1 min.

After the PCR products were cleaned, their DNA content was quantified using the Qubit method described above. The concentration in nM was calculated using Equation S26 [59]. The average library size of 390 bp was used. The pooled sample was diluted to a concentration of 4 nM using Resuspension Buffer. The sample was stored at $-20\,^{\circ}$C.

$$\frac{\text{concentration in } \frac{ng}{\mu L}}{660\frac{g}{mol} \times \text{average library size}} \times 10^6 = \text{concentration in nM} \tag{S26}$$

##### 8.2.3 MiSeq sequencing

Illumina's 16S Library preparation protocol [59] was used with some modifications from the Earth Microbiome Project's protocol for the final denaturing and sequencing steps. Paired end sequencing with 150 bp using MiSeq reagent kit V2 for 300 cycles (catalog number: MS-102-2002) was performed. Fresh 0.2 N NaOH was prepared immediately prior to the denaturation steps. 5 $\mu$L of the pooled DNA library was mixed with 5 $\mu$L 0.2 N NaOH. The mixture was vortexed briefly to and centrifuged at about 280 g for 1 min. The mixture was then incubated at room temperature for 5 min to denature the DNA. 990 $\mu$L of pre-chilled HT1 solution was added to the denatured DNA. The resulting 20 pM denatured DNA library was placed on ice.

PhiX control from Illumina (Catalog number: FC-110-3001) was used to make the sample more complex. The 10 nM library was diluted to 4 nM using resuspension buffer. It was denatured by mixing 5 $\mu$L of the 4nM PhiX library with an equal volume of 0.2 N NaOH. After vortexing, the mixture was incubated for 5 min at room temperature to allow denaturation. Then, 990 $\mu$L of pre-chilled HT1 buffer was added to create a 20 pM denatured phiX library.

Both the phiX and the DNA libraries were diluted to 8 pM by mixing 360 $\mu$L of HT1 solution with 240 $\mu$L of the libraries in separate microcentrifuge tubes. 5% phiX was used in the final library by mixing 30 $\mu$L of the 8 pM denatured phiX library with 570 $\mu$L of the 8 pM denatured DNA library. The mixture was placed on a heat block pre-heated to 96 C for 2 min. Then, the mixture was inverted a few times to ensure mixing and placed on ice for 5 min.

Meanwhile, the thawed reagent cartridge was prepared for sequencing by gently flipping up-down for about 10 times and tapping on a table to ensure all reagents were collected at the bottom of the wells. Wells 12, 13 and 14 were pierced using a pipette tip. 3.4 $\mu$L of 100 $\mu$M index sequencing primer was added to well 13, 3.4 $\mu$L of 100 $\mu$M read 1 sequencing primer was added to well 12, and 3.4 $\mu$L of 100 $\mu$M read 2 sequencing primer was added to well 14. The contents of the well were mixed using pasteur pipettes. The sample on ice was added to well marked "Load Sample." The cartridge was then loaded to the MiSeq and a new .csv file was made to incorporate the changes made by the Earth Microbiome Project protocol. The sequencing was then started.

#### 8.3 Data processing

Once the sequencing run was completed, the data was converted to fastq format on the sequencing machine using the MiSeqReporter service. The data was then transferred for further analysis. The run yielded 5.65 Gbp of data, of which 89% had Qscore greater than 30. Qiime2's [60] "Moving Pictures Tutorial" [61] was used as a basis to demultiplex the paired end reads and export them to fastq format. This data was then imported to R, where the DADA2 [62] pipeline v1.6 [63] was used to filter, trim, denoise, remove chimeras, and merge the paired end reads. The normalization of the different communities was checked at the beginning of the pipeline, in terms of number of total reads per sample, as shown in Figure S23. The SILVA database [64] was used to assign phylogenetic information. From here, a sequence table containing the number of reads of each sequence per sample and a table of phylogenetic information of each sequence were obtained. This information is imported to python for further analysis. 992 exact sequence variants were identified among all the samples.

### 9 16S sequence data analysis

The sequence table and the associated phylogenetic table was imported into Python. First, any sequence not associated with a Kingdom was removed. Next, from the control sample of *C. reinhardtii*, the 16S sequence corresponding to its chloroplast was found. This sequence was then removed from all other samples. The number of reads were then converted to relative abundance for each community by dividing the number of reads for each sequence by the total number of reads for that community.

Since the reads were converted to relative abundances, each community could be viewed as a normalized probability distribution of the sequences it contains. To quantify the similarities and differences between different communities, the Jensen Shannon divergence metric was used. This metric is better than other Shannon entropy based measures because it is bounded, has the capability to be weighted and is symmetric [65]. In general, the Jensen Shannon divergence between two normalized probability distributions, $X$ and $Y$, is given by Equation S27, where H is the Shannon entropy of the probability distribution and $\pi_1$ and $\pi_2$ are the weights for the two distributions.

$$J_{X,Y} = H(\pi_1 X + \pi_2 Y) - \pi_1 H(X) - \pi_2 H(Y) \tag{S27}$$

If the distribution $X$ is given by $X = \{x_i\}$, where $x_i$ represent the normalized probability of finding the value $x_i$ in the probability distribution $X$, the entropy $H$ of the probability distribution $X$ is defined by Equation S28.

$$H(X) = -\sum x_i \log x_i \tag{S28}$$

Here, we set the weights to be $\pi_1 = \pi_2 = \frac{1}{2}$. Using Equation S27, we can now define the Jensen Shannon divergence between the relative abundance composition of CES. Let $A_i^d$ and $B_i^d$

denote the normalized distributions of relative abundances, which are equivalent to probability distributions, for CES derived from soil samples A and B respectively, where $i \in \{1,2,3,4\}$ represents the four communities derived from each soil sample, and $d \in \{1,2,3,4\}$ represents the the four dilution rounds.

The intra-CES A Jensen Shannon divergences plotted in Figure S13 are obtained by computing Equation S29 for all four $A_k$ and for all six pairs of $\{d_l, d_m\}$, resulting in 24 different combinations. Similarly, intra-CES B Jensen Shannon divergences are computed for communities derived from soil sample B, by replacing $A$ with $B$ in Equation S29. These divergences indicate how similar each CES is to itself over the four rounds of dilutions.

$$J_{A_k}^{d_l,d_m} = H(\frac{A_k^{d_l} + A_k^{d_m}}{2}) - 0.5H(A_k^{d_l}) - 0.5H(A_k^{dm}) \tag{S29}$$

Inter-CES A Jensen Shannon divergences in Figure S13 are calculated by Equation S30 for all unique pairs of $\{A_i, A_j\}$ between all pairs of dilution rounds, $\{d_l, d_m\}$, resulting in 96 pairs. Similarly, Inter-CES B Jensen Shannon divergences are computed using Equation S30, by replacing $A$ with $B$. These divergences indicate how similar each CES is to other CES derived from the same soil sample.

$$J_{A_iA_j}^{d_l,d_m} = H(\frac{A_i^{d_l} + A_j^{d_m}}{2}) - 0.5H(A_i^{d_l}) - 0.5H(A_j^{d_m}), i \neq j \tag{S30}$$

The Jensen Shannon divergences of relative abundances between the communities derived from the two soil samples, plotted in Figure S15 are computed in Equation S31 for each dilution round $d$ for every unique $\{A_i, B_j\}$ pairs, resulting in 16 divergences at each dilution round. These divergences indicate how similar CES derived from soil sample A are to CES derived from soil sample B at each dilution round.

$$J_{A_iB_j}^{d} = H(\frac{A_i^{d} + B_j^{d}}{2}) - 0.5H(A_i^{d}) - 0.5H(B_j^{d}) \tag{S31}$$

The Jensen Shannon divergences of the relative abundances were then computed between all the 32 communities. These distances were embedded using Multi-dimensional scaling [66] to aid in visualization. Scikit-learn's [67] "mds" method was used to embed the data. The results are shown in Figure 3B. The method minimizes stress S, which is an objective function that measures how accurately the embedding describes the measured distances between communities. If $J_{A_i,B_j}$ is the Jensen Shannon divergence between communities $A_i$ and $B_j$, and $S_{A_i,B_j}$ is the distance in the embedded coordinate system, the stress S is given by

$$S = \sum J_{A_iB_j} - S_{A_i,B_j} \tag{S32}$$

where $J_{A_iB_j}$ is the JSD between communities $A$ and $B$ and $S_{A_i,B_j}$ is the distance in the embedded coordinate system. The summation runs over all pairs of CES. This stress depends on the number of embedding dimensions. Figure S18 shows the stress as a function of the number embedding dimensions. At two dimensions, since the curve begins to plateau, and the stress is very close to the minimum stress, two embedding dimensions are used in Figure 3B of the main text.

#### 9.1 Aitchison's distance

For compositional data like the 16S data presented here, it is recommended [68] to use Aitchison's distance to quantify differences between communities. Aitchinson's distance accounts for the compositional nature of the data and avoids artifacts resulting from the fact that the data lie on a simplex. While the JSD metric is robust to this constraint (since it measures distances between normalized distributions) we checked that using Aitchinson's distances did not alter our results regarding community taxonomic structure. We used the zCompositions tool in R [69] to replace zeros using the cmultRepl function, which replaces zero counts using a Bayesian-multiplicative replacement [70], using a geometric prior and scales the non-zero counts. This results in a corrected relative abundance table for all CES. This table was ported back to Python and the center log-ratio (clr) was found for the now non-zero relative abundances for each CES. Then, the Euclidean distance was calculated between every pair of CES [71], giving the Aitchison's distance. This distance is used as a metric in Figure S14 which validates our analysis using Jensen Shannon Divergence.

#### 9.2 OTU clustering

OTU clustering was performed to ensure that the results hold when similar ESVs were grouped, as shown in Figure S16. For such clustering, the "dada2 denoise-paired" function was called within the Qiime2 pipeline after demultiplexing the sequences. The vsearch [72] plugin was called through Qiime2, to cluster these de-noised reads. This was repeated for different similarity thresholds and the results were imported as feature tables. The divergences between communities using different similarity measures to define OTUs were computed as explained above. The results are shown in Figure S16. For a phylogenetic tree of the ESVs observed in all CES at all rounds see Figure S17.

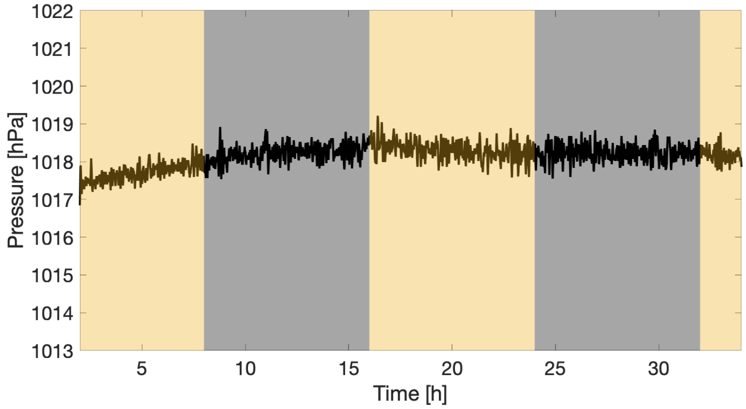

**Figure S1: Pressure measurement control experiment with a vial containing** $20\,\mathrm{mL}$ **of water only.** $8\,\mathrm{h}$-$8\,\mathrm{h}$ light-dark cycles with an intensity of $150\,\mathrm{\mu mol\,m^{-2}\,s^{-1}}$ were applied while the vial was held under active temperature control as described in the Methods. Note that the pressure does not change in response to illumination. The time series has not been smoothed.

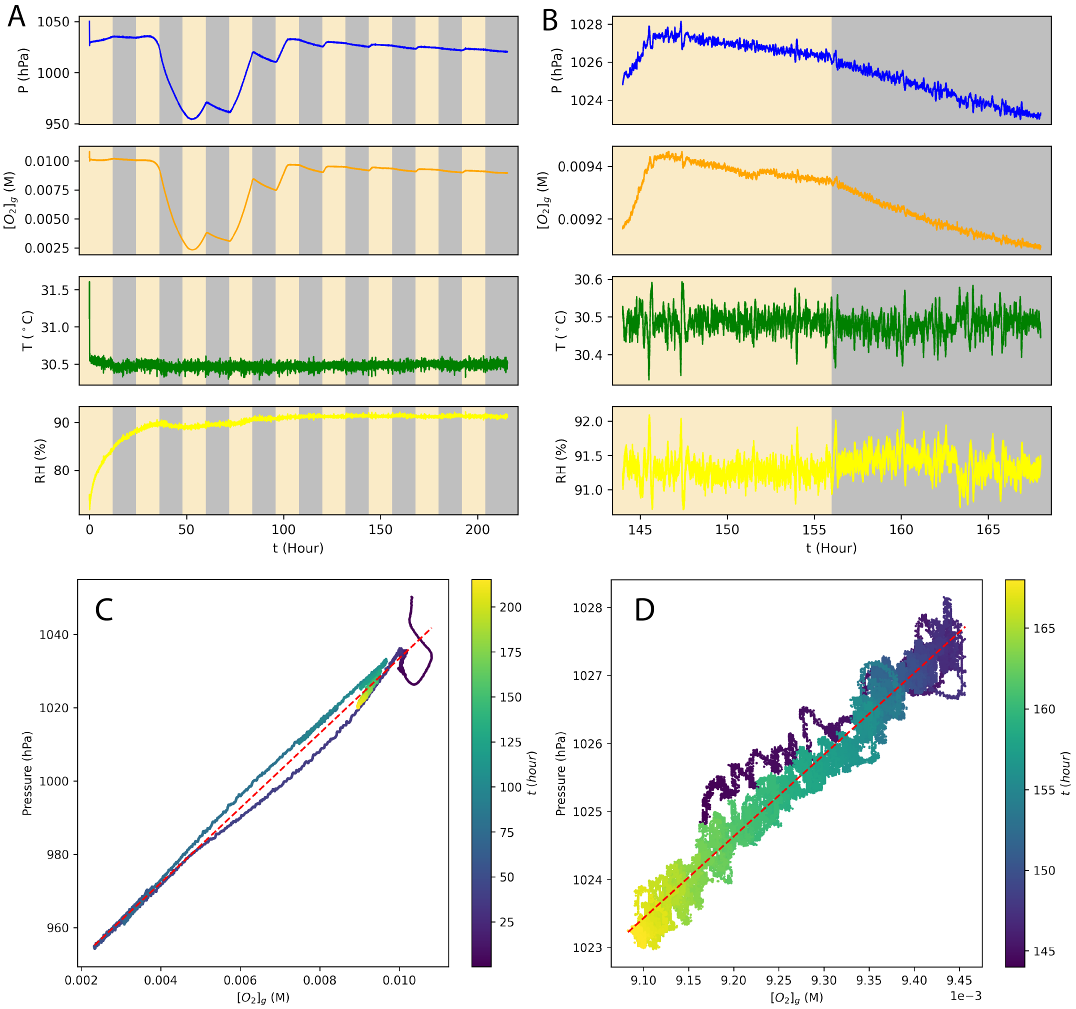

**Figure S2: Control experiment with both pressure and $O_2$ concentration measurements.** (A) Pressure (hPa), temperature (°C), relative humidity (%) (all measured via BME280 sensor) and gaseous $O_2$ molarity concentration (M) (measured via Presens sensor) as a function of time for a CES subjected to 12h-12h light-dark cycles. (B) Data in (A) on 6th light-dark cycle. (C) Pressure varies with $[O_2]_g$ linearly (slope=$10206.5 hPa/M$, $r^2 = 0.994$). Color indicates time in hours. (D) Data in (C) on 6th day-dark cycle (slope=$12024.2 hPa/M$, $r^2 = 0.962$).

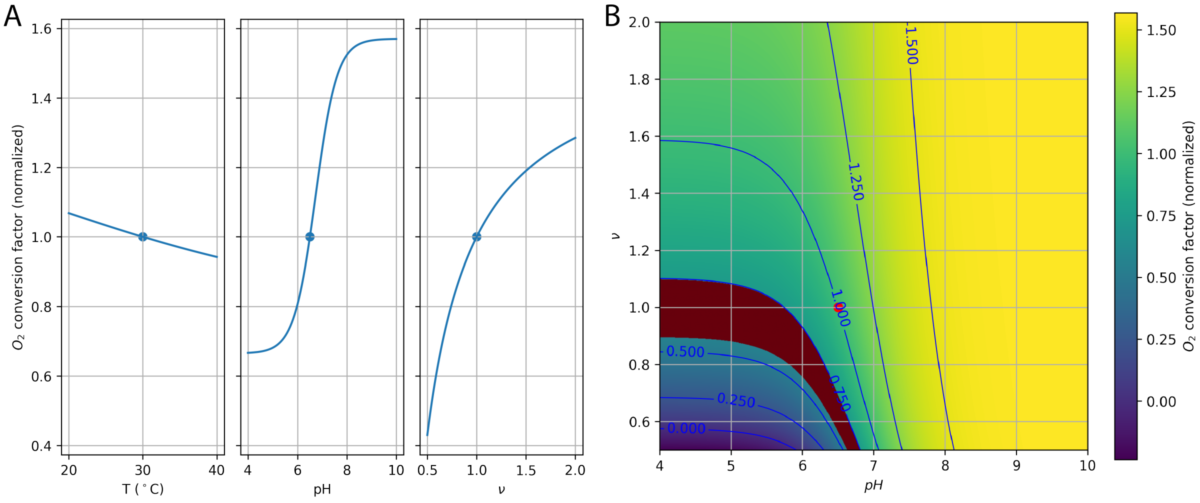

**Figure S3: Dependencies of the $O_2$ conversion factor on $pH$ and $\nu$.**(A) The $O_2$ conversion factor ($\Delta P/\Delta[(O_2)]_g$, Equation S14) weakly depends on temperature but strongly varies with $pH$ and $\nu$. In each panel, conversion factors are calculated with one of $(T, pH, \nu)$ perturbed from the default parameters ($T = 30\,°\mathrm{C}$, $pH = 6.5$, $\nu = 1$; blue dots) while the other two variables are held fixed, and then normalized by dividing the reference conversion factor at $T = 30\,°\mathrm{C}$, $pH = 6.5$, $\nu = 1$. (B) The $O_2$ conversion factor (normalized) in the $(pH, \nu)$ parameter space. The red shaded region corresponds to the experimentally measured conversion factor values. The red dot indicates the default parameters.

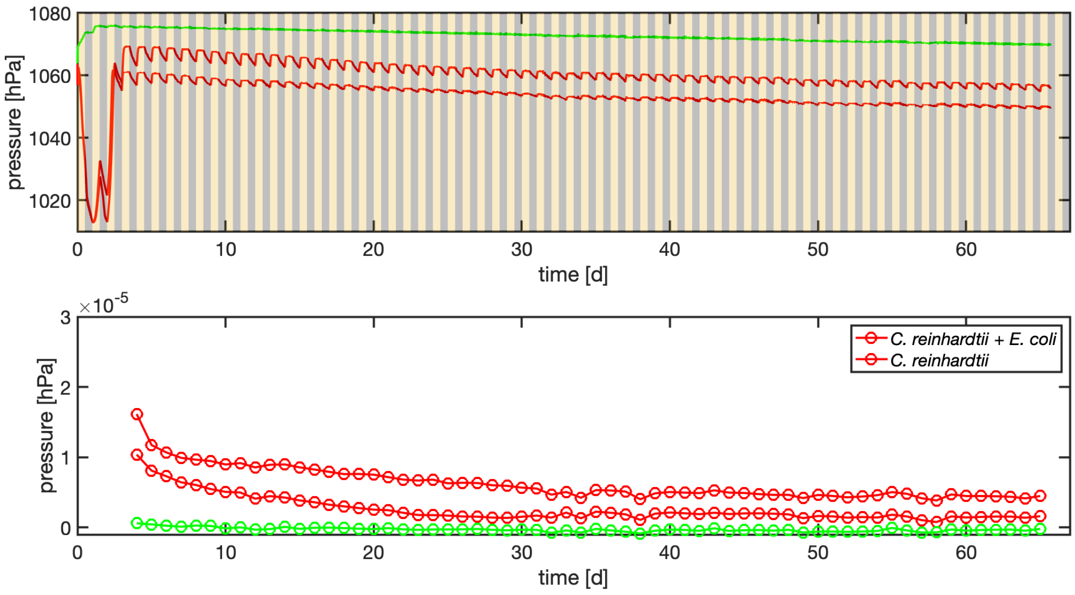

**Figure S4: Raw data for *C. reinhardtii* and *C. reinhardtii  E. coli* controls.**(top panel) Time series of pressure in time for CES containing either *C. reinhardtii* alone (green) or *C. reinhardtii* + *E. coli* (red). (bottom panel) Carbon cycling rate for the three synthetic CES shown above. The rates of the two *C. reinhardtii* + *E. coli* replicates are averaged and shown in Figure 2 of the main text.

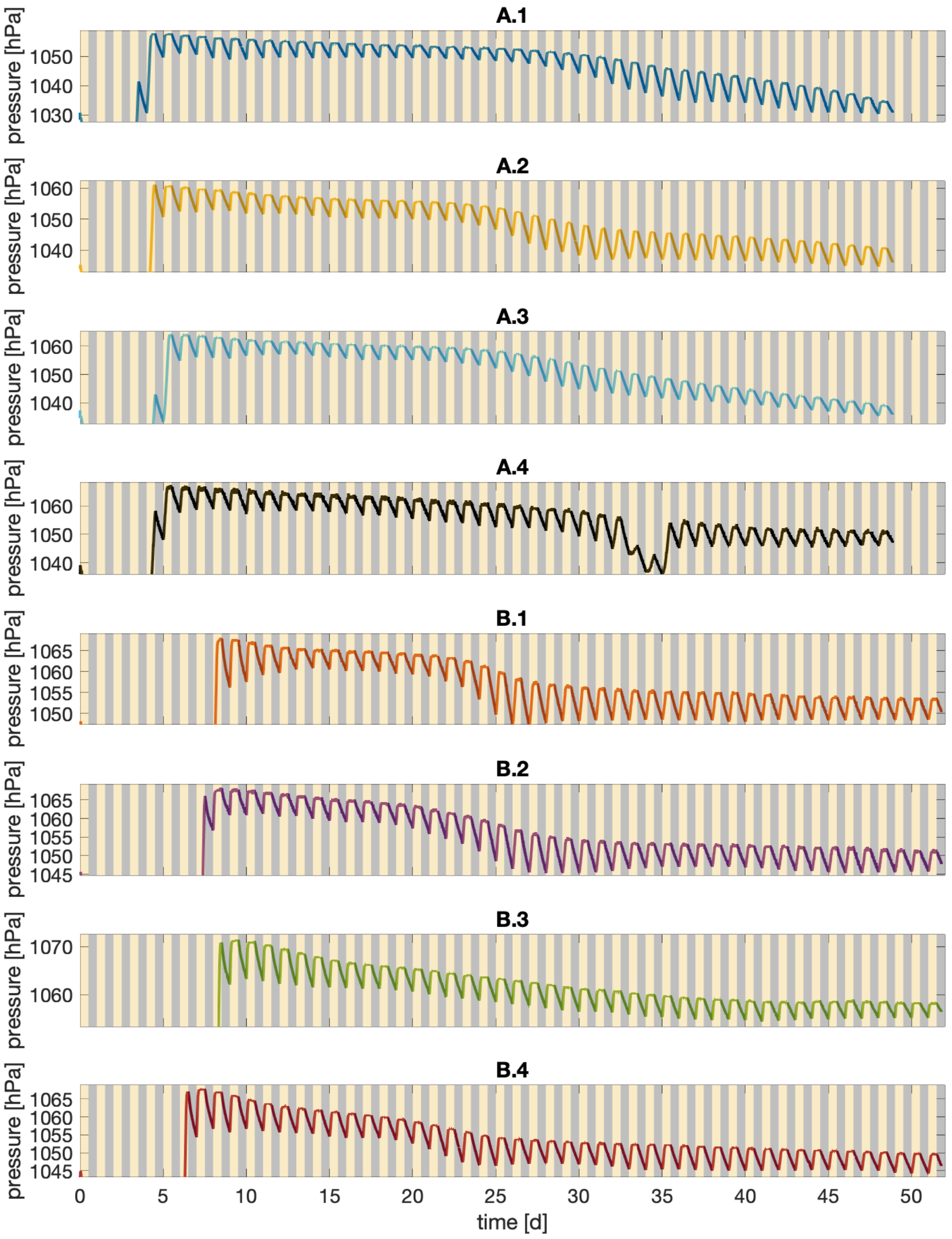

**Figure S5: Pressure data for all eight CES during the first round of closure.** Data identical to that shown in panel (A) of Figure 2 of the main text. Soil sample and CES number are given in the titles of each panel and correspond to the legend in Figure 2 of the main text. Axes limits are set to omit the initial transient period for clarity.

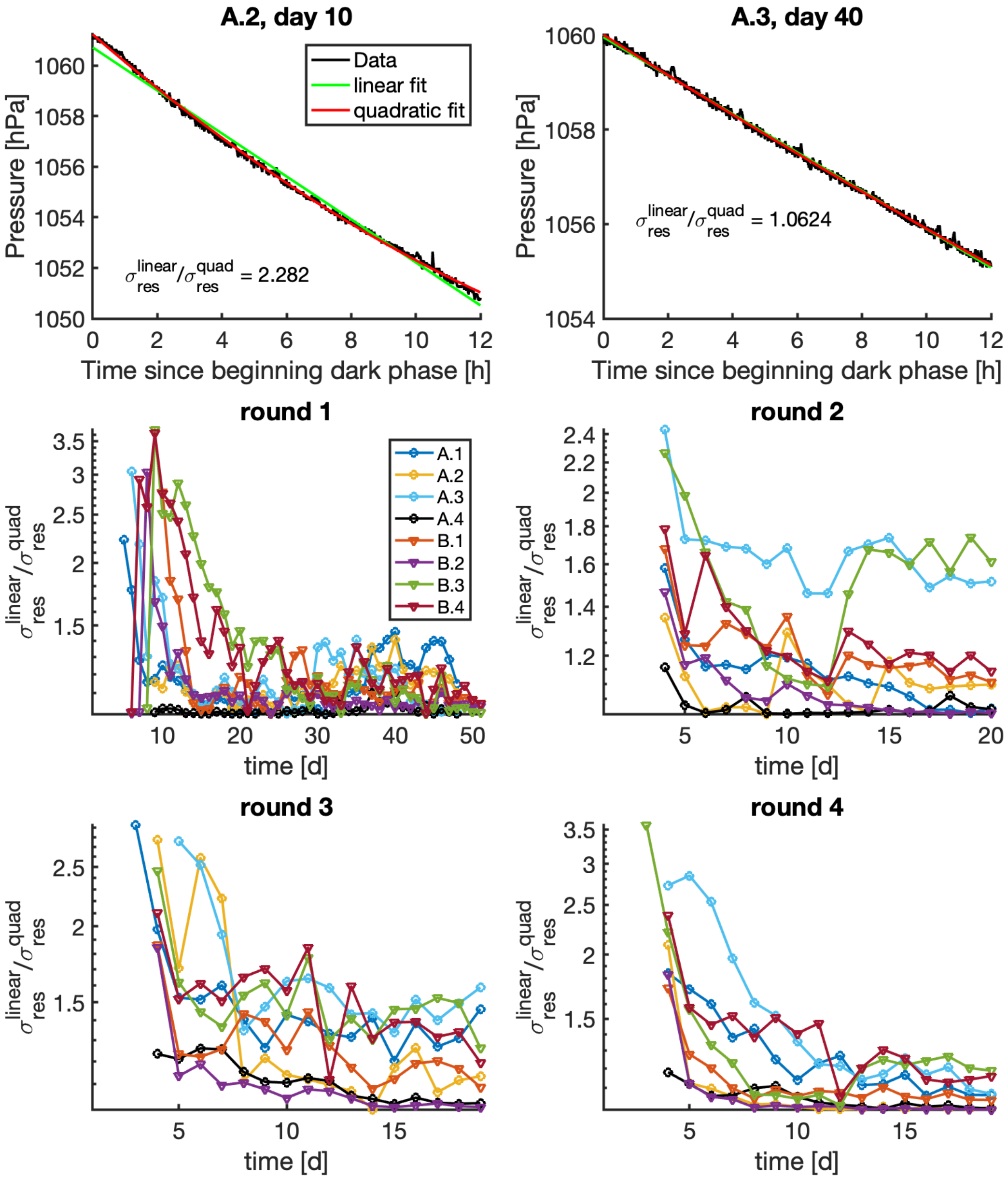

**Figure S6: Quantifying constancy of dark phase respiration rates.** (top two panels) For all CES during all four rounds of closure we extract the pressure data for each dark phase. The top two panels show two such examples from CES as shown in the panel titles. For each dark phase a linear (green) and quadratic (red) polynomial is fit to the data by ordinary least squares and the residual is computed. We then compute the standard deviation of the residual for each model $\sigma_{res}^{linear}$ and $\sigma_{res}^{quad}$ and the ratio of these two quantities as shown. (bottom four panels) Show the ratio $\sigma_{res}^{linear}/\sigma_{res}^{quad}$ as a function of time for all 8 CES during all four rounds of dilution as shown in the panel titles. The legend from the first round applies to all four panels and corresponds to Figure 2 of the main text.

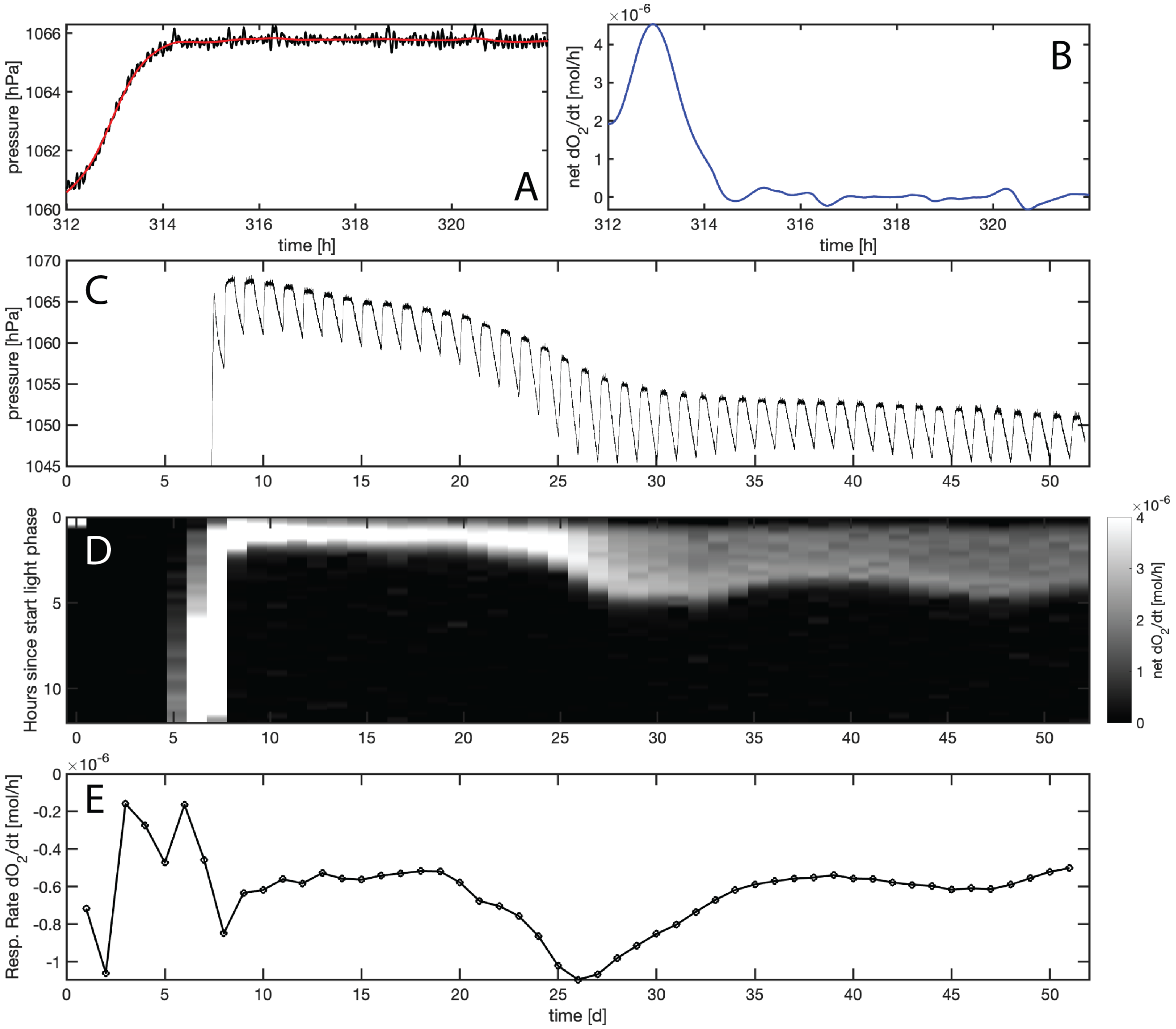

**Figure S7: Rates of photosynthesis and respiration during round 1 in CES B.2.** (A) Pressure in time during a single light-phase for CES B.2 on day 13 (Figure 2, main text). Black line shows data and the red line is a smoothing spline fit to the data. (B) The time derivative of the smoothing spline fit from (A). Units are converted from pressure to $O_2$ rates assuming $\nu = 1$ and pH 6.5. Net oxygen production rates means that respiration is not accounted for in the calculation. (C) Pressure in time for round 1 of CES B.2 (as in Figure 2, main text). (D) Net $O_2$ production rates for all light phases during round 1 for CES B.2. Columns show time since the beginning of each light phase (y-axis) in time (x-axis). Heat map is net $O_2$ production rate as shown in the color bar to the right. (E) Estimated rate of consumption of $O_2$ by respiration during the corresponding dark phases over the course round 1 CES B.2. Note the negative values indicating consumption.

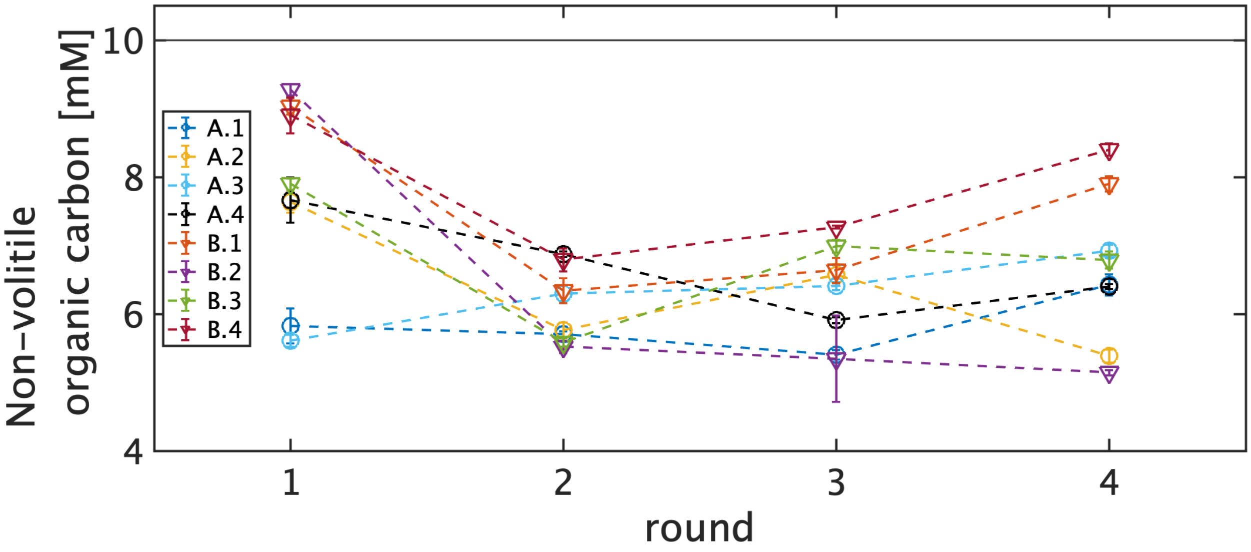

**Figure S8: Total organic carbon for each CES at the end of each round.** For discussion of the measurements see Section 7.4. The gray line indicates the concentration of organic carbon supplied at the initiation of each CES by the media (Table S5). CES are identified in the legend.

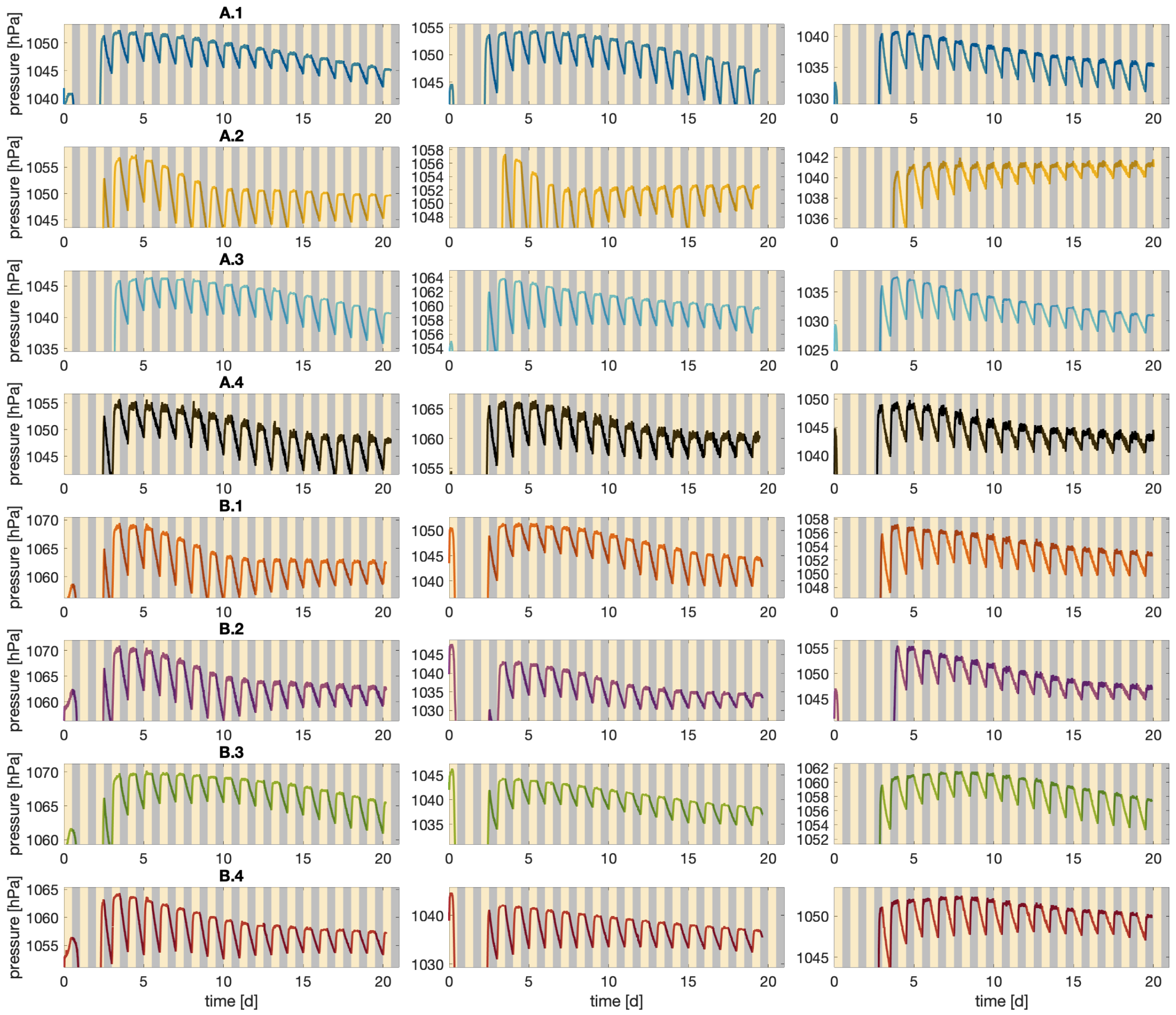

**Figure S9: Pressure data for all eight CES during the three enrichment steps shown in panels (C-E) of Figure 2.** Each row corresponds to one CES identified in the title of the panel on the left. Colors correspond to legend in Figure 2 of the main text. Axes limits are set to omit the initial transient period for clarity.

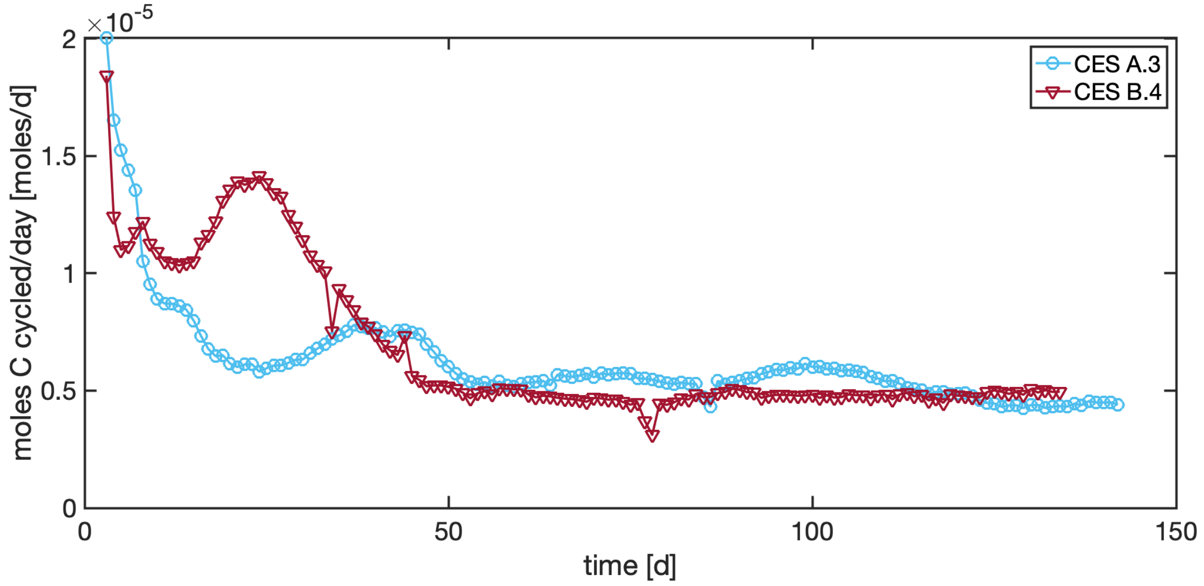

**Figure S10: Long-term carbon cycling in two CES.** Carbon cycling rates in two CES which were diluted and sealed again at the end of round 4.

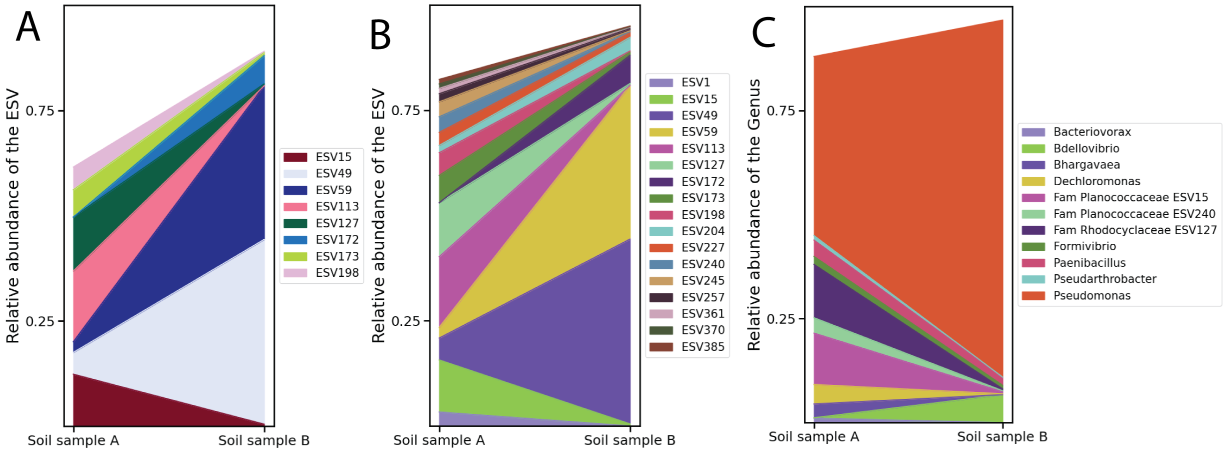

**Figure S11: Composition of the two initial soil samples after treatment with drugs.** (A) shows the ESV level composition of the two soil samples used to start the experiment, after treatment with drugs to remove fungi and other eukaryotes. Only those ESVs with a relative abundance of 5 % or more in either samples are plotted. The colors of the ESVs are the same as in Figure 3A. (B) shows the ESV level composition of the two soil samples with a cutoff of 1 %. (C) shows the Genus level abundances of the two soil samples. ESVs with common Genus labels are combined. If the Genus is not assigned, the name of the next higher taxonomic rank is assigned, with the name of the rank as a prefix, and the ESV label is the suffix. Here, "Fam" in the legend denotes the taxonomic rank Family. Only those genera that have a relative abundance of 5% or more in at least one soil sample are included. For a complete list of taxa in each sample see Supplementary Data 1. The Jensen Shannon divergence between soil sample A and the CES derived from it at the end of the first dilution is 0.68 ± 0.014, and for soil sample B is 0.69 ± 0.001. Only ESV15 and ESV1 are present in the CES at more than 5 % abundance.

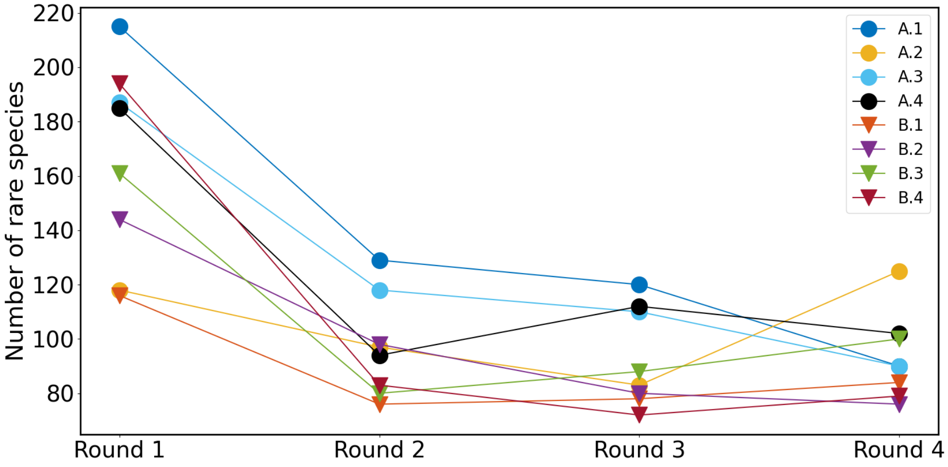

**Figure S12: Number of rare taxa in CES.** The number of rare taxa, defined as the taxa with a relative abundance of less than 5%, decreases across all communities as a function of dilution rounds.

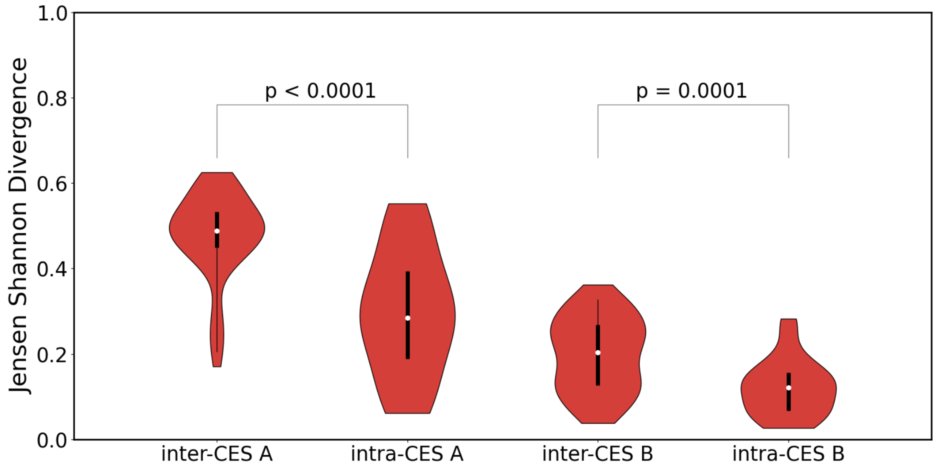

**Figure S13: The distribution of Jensen Shannon divergences between the CES, based on the relative abundances of ESVs.** The relative abundance is obtained by the 16S sequences as described in Section 8.2.3. The Jensen Shannon divergence is then calculated as in Equation S27. The intra-CES Jensen Shannon divergences were calculated using Equation S29 between each community at different dilution rounds e.g. for CES A.1 between rounds 1, 2, 3 and 4. There are 6 intra-CES divergences for each community and therefore 24 for each soil sample. The inter-CES Jensen Shannon divergence is calculated by Equation S30 between different CES communities e.g. between CES A.1 at round 1, and all other CES of sample A at all 4 rounds. This results in a total of 96 unique divergences for each soil sample. Violin plots compare the distributions of inter- and intra-CES divergences. The intra-CES distribution has lower median values than the inter-CES distributions for both soil samples. The p-values are calculated by bootstrapping to test for the null hypothesis that the median of the distributions of the Jensen Shannon divergences for inter and intra CES are the same. The low p-values for both soil samples allows us to reject the null hypothesis.

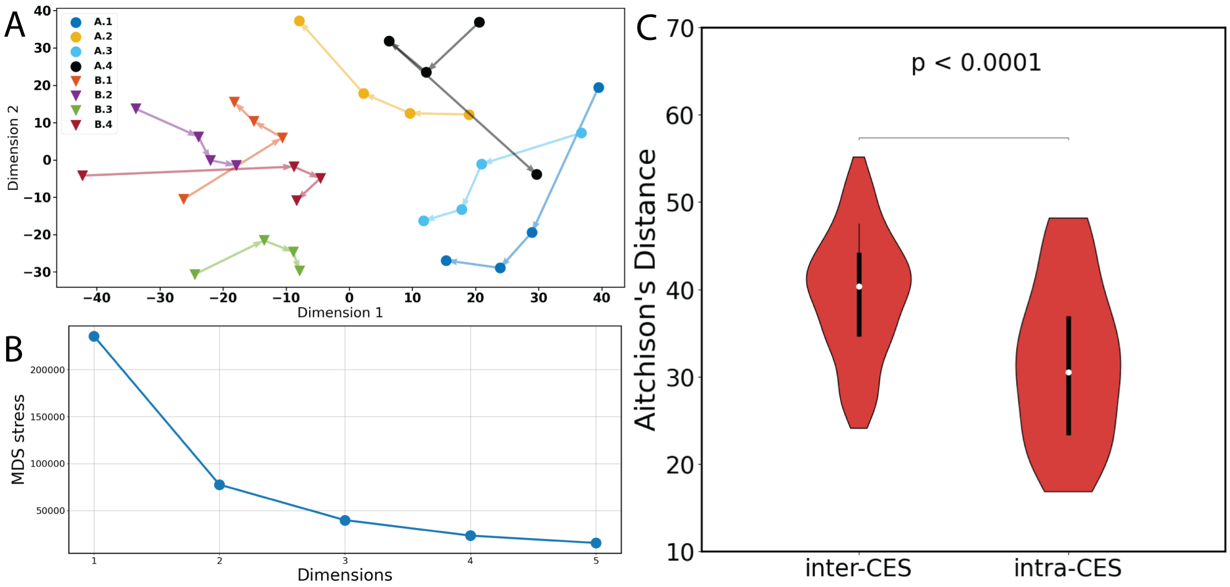

**Figure S14: Taxonomic differences hold when Aitchison's distance is used as a metric.** The Aitchison's distances were calculated as described in section 9.1. (A) shows the 2 dimensional Multi Dimensional scaling embedding of the Aitchison's distances. (B) shows the stress of embedding as a function of the number of embedding dimensions. (C) The distances for inter and intra-CES for the two soil types are combined here resulting in 192 distancess for the inter-CES samples and 48 for the intra-CES samples The p-values are calculated by bootstrapping for the null hypothesis that the inter and intra CES distances have the same median. The low p-values refute the null hypothesis.

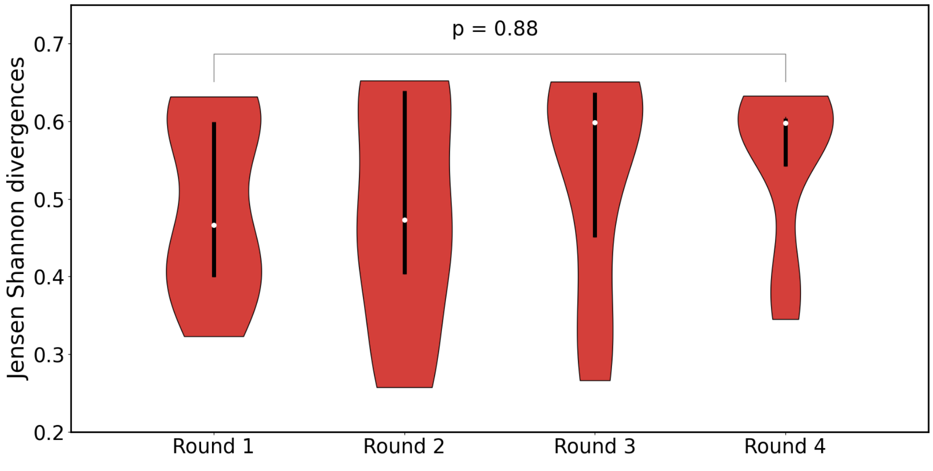

**Figure S15: Jensen Shannon divergences of relative abundances of ESVs between CES derived from the two soil types.** The Jensen Shannon divergences of the relative abundances are calculated using Equation S31 between CES belonging to different soil types for each dilution round, e.g., the divergence between A.1 and B.1, B.2, B.3 and B.4. There are 16 such divergences for each dilution round. There is no decline in the median divergence over dilutions, as shown by the p-value calculated by bootstrapping for the null hypothesis that the median of the distributions of Jensen Shannon divergences between the two soil samples is the same for round 1 and round 4. The high p-value indicates the null hypothesis cannot be ruled out.

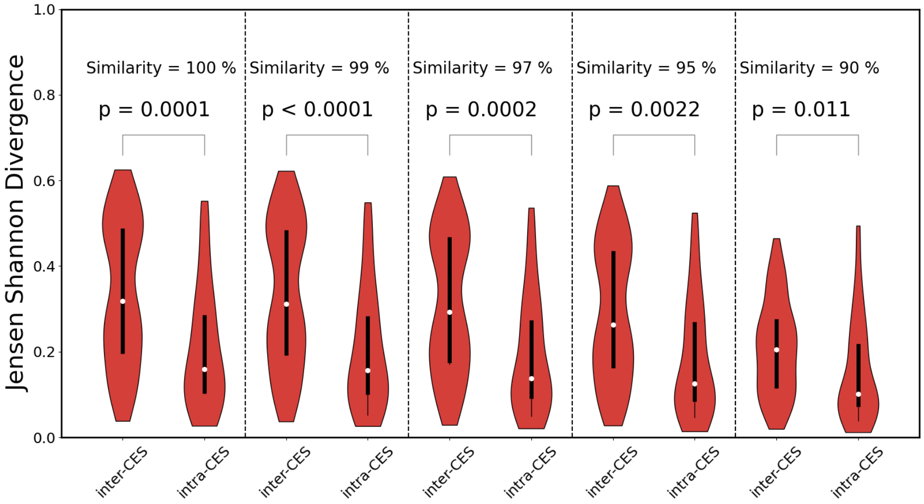

**Figure S16: Taxonomic differences are preserved on coarse-graining the 16S sequences.** The Jensen Shannon divergences for inter and intra-CES were calculated as described in Figure S13. The divergences for the two soil types are combined here resulting in 192 divergences for the inter-CES samples and 48 for the intra-CES samples at each similarity level. This was then performed at various levels of coarse-graining the 16S sequence similarity, indicated by the similarity percentage above each pair of inter and intra-CES divergences. The p-values are calculated by bootstrapping for the null hypothesis that the inter and intra CES divergences have the same median. The low p-values refute the null hypothesis.

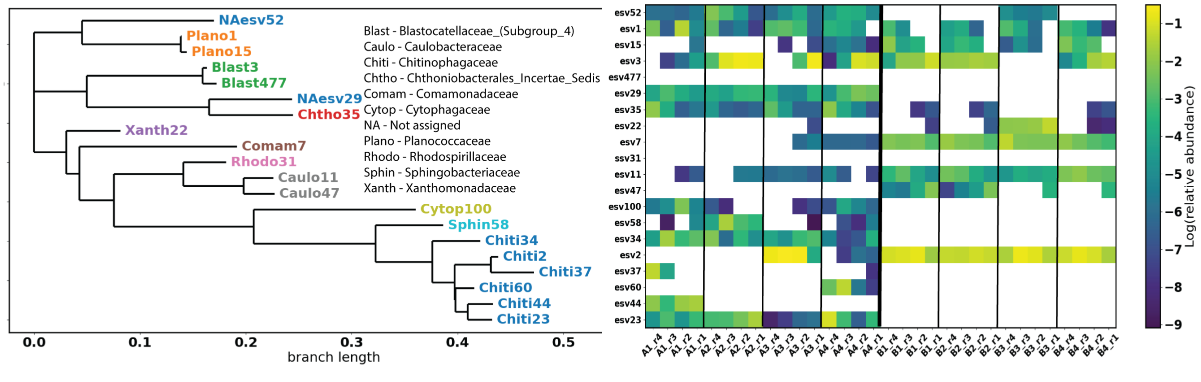

**Figure S17: Phylogenetic tree of ESVs detected in CES.** For ESVs present at a relative abundance of at least 10 % in any time point in any CES a phylogenetic tree was constructed using the SILVA alignment and classification tree service (`https://www.arb-silva.de/aligner/`) using the FastTree algorithm. For each branch the family identity is shown by the color. Branch length is in units of substitutions per site. The heat map at the right shows the log relative abundance of each taxa where white indicates the taxa is not observed. Labels across the bottom indicate the CES and round.

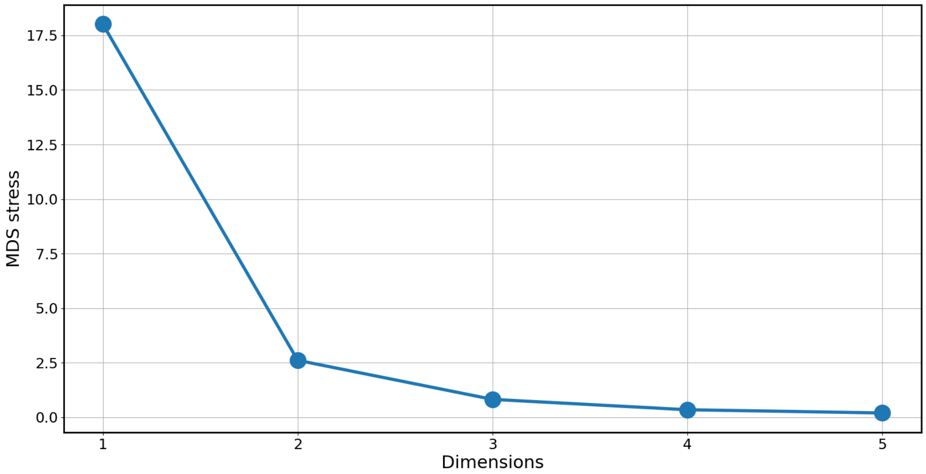

**Figure S18: The stress of the Multi Dimensional Scaling (MDS) method for embedding Jensen Shannon divergences between CES based on the relative abundances of Exact Sequence Variants(ESVs), as a function of number of embedding dimensions.** The stress (Equation S32) reported by the MDS method used to embed the Jensen-Shannon divergences between the CES is plotted on the y-axis. The divergence is calculated based on the relative abundances of the ESVs. The x-axis shows the number of spatial dimensions used for the embedding.

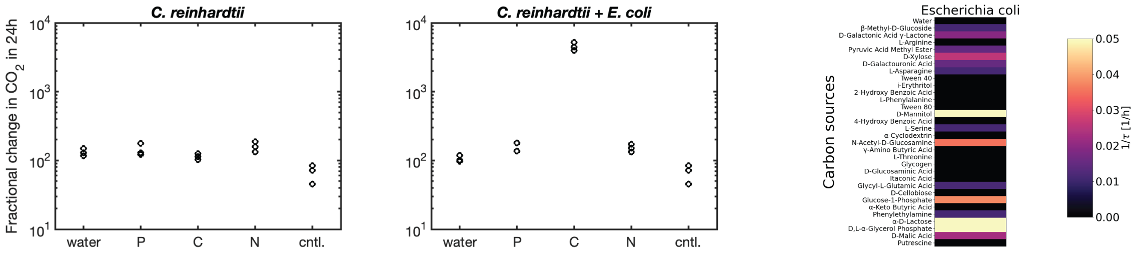

**Figure S19: Microresp and EcoPlate measurements for control synthetic CES** (left two panels) Microresp measurements for CES comprised of only only *C. reinhardtii* or *C. reinhardtii* + *E. coli*. Compare to Figure S21. (right) Ecoplate data for *E. coli* alone. The heat map is identical to Figure 3 of the main text. Values are averages across three replicates for each carbon source.

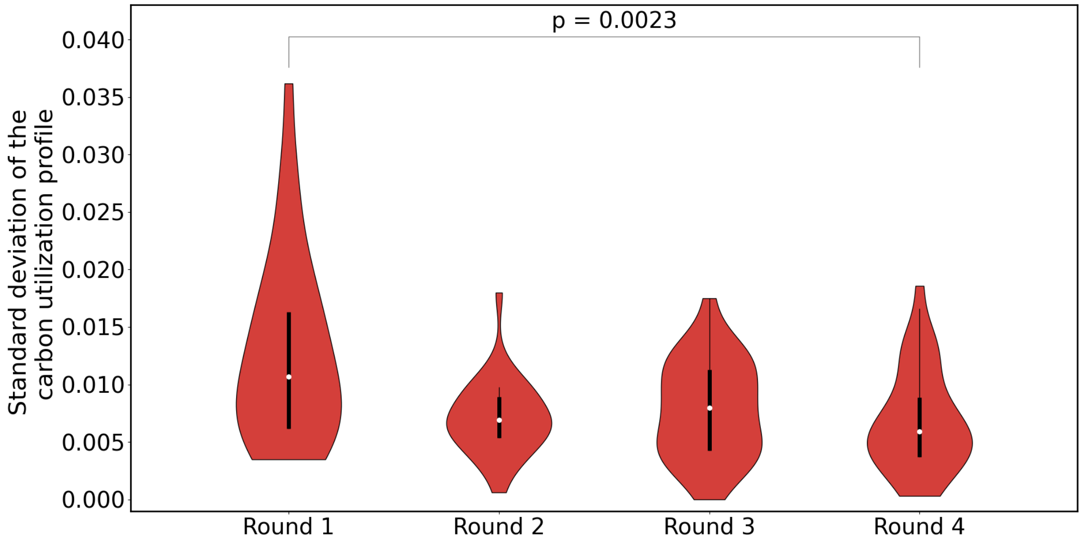

**Figure S20: Standard deviation in the carbon utilization profile of the CES converges over dilution rounds.** The carbon utilization profile of the CES is defined as the consumption rate of various carbon compounds in a Biolog EcoPlate. Here, the standard deviation in the consumption rate is calculated for each carbon source across all 8 CES, each with three replicates, resulting in 32 standard deviations, as shown in Equation S23. The violin plots show the distribution of these standard deviations for each round of enrichment. The p-value is calculated by bootstrapping to test for the null hypothesis that the medians of the distributions of the standard deviations of dilution round 1 and dilution round 4 are the same. The low p-value refutes the null hypothesis.

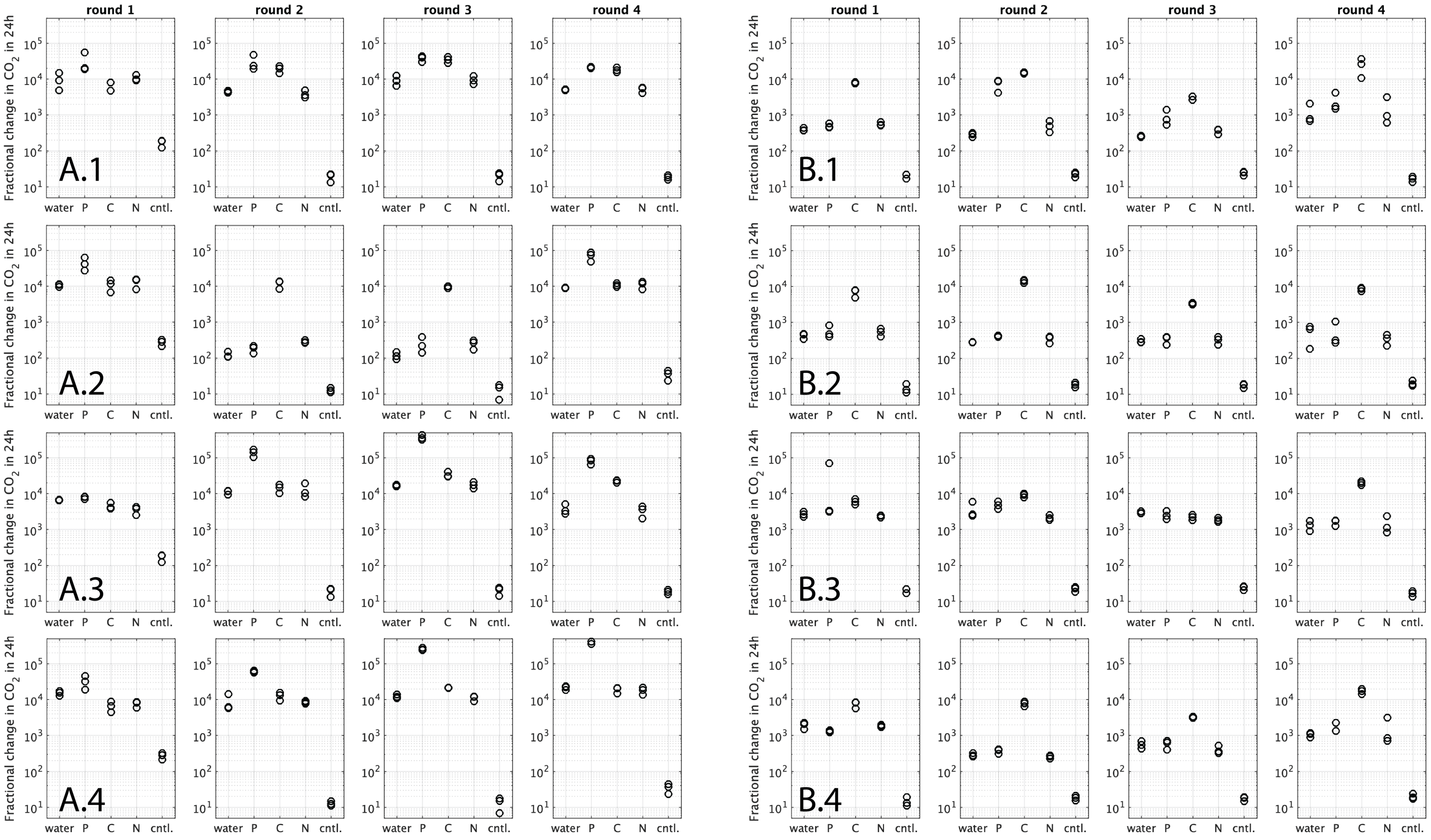

**Figure S21: Nutrients limiting respiration.** Measurements of respiration at the end of each round for all CES. CES from sample A on the left and B on the right. Each panel shows fractional change in $CO_2$ (Equation S25) produced in 24 h period for an sample of each CES. In each column, the CES sample is amended with water, P (phosphate, $KH_2PO_4/K_2HPO_4$), C (carbon, glucose), N (nitrogen, $NH_4Cl$). The control condition ('cntl') contains only water (no cells). Each condition is assayed in triplicate.

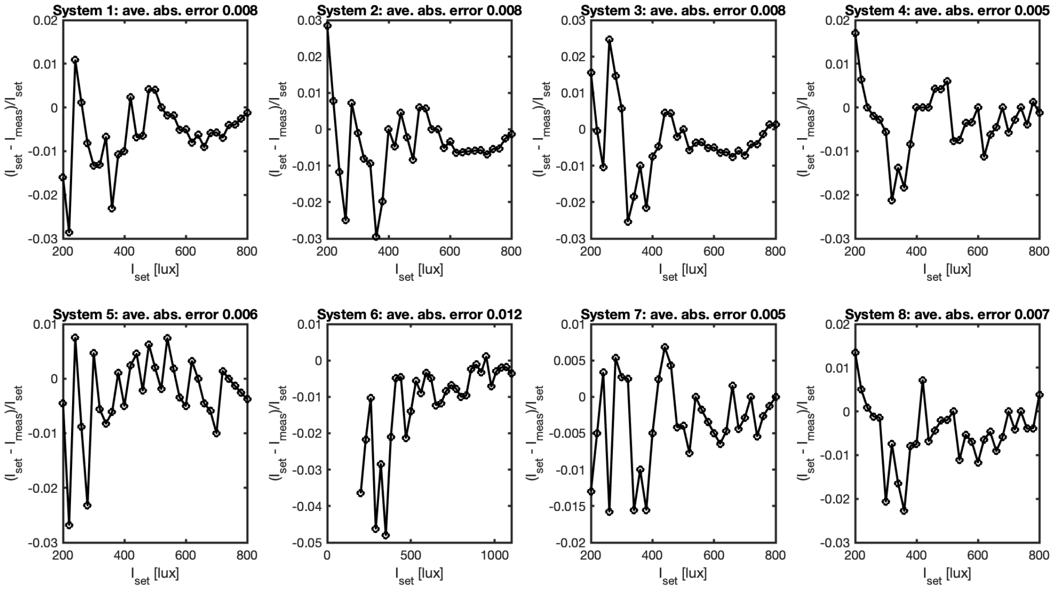

**Figure S22: Calibration of LED illumination in custom culturing devices.** Independent calibration were performed for all 8 culture devices. Plots show set LED intensity verses fractional error as denoted on the y-axis for each panel. The 'ave. abs. error' in each panel denotes the mean of the absolute value of all points in each panel.

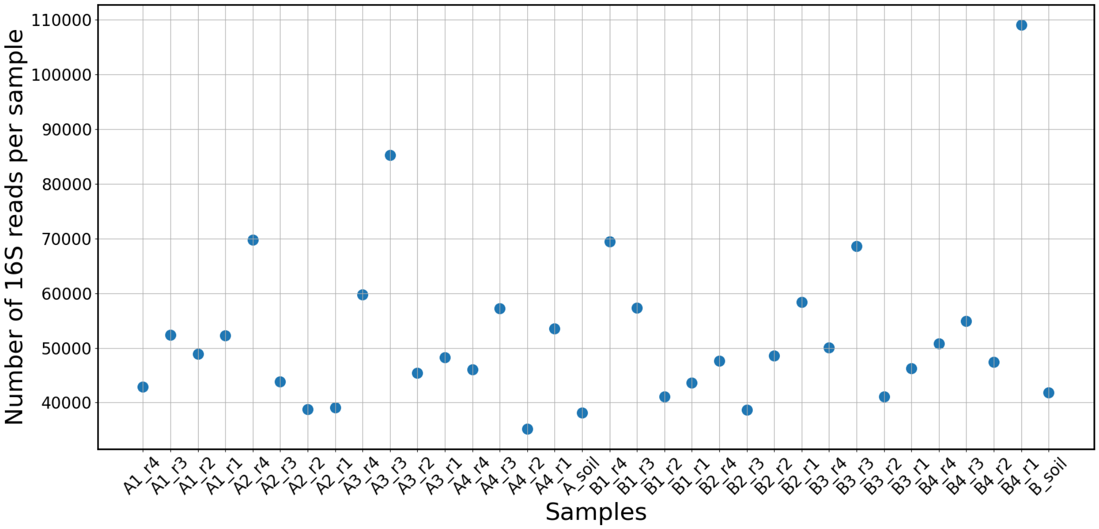

**Figure S23: The number of reads obtained per sample after processing them through the DADA2 pipeline.** In the sample names, r1, r2, r3, r4 correspond to dilution rounds 1, 2, 3 and 4 respectively, and "A_soil" and "B_soil" correspond to the initial soil samples used to start the cultures.

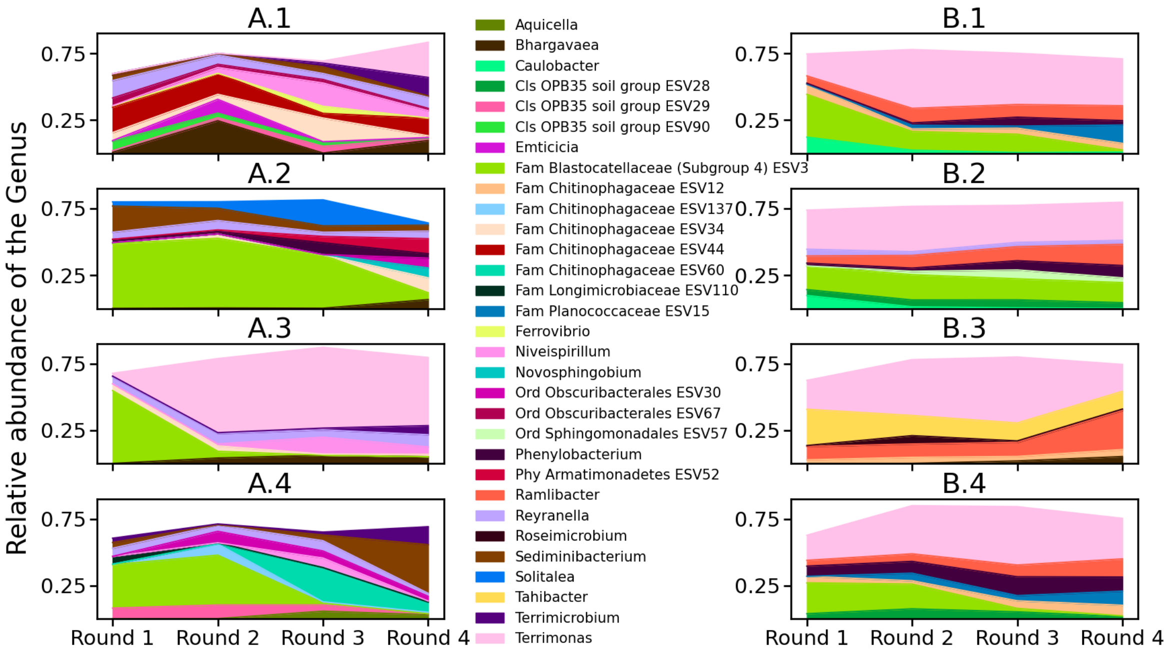

**Figure S24: Time series of the genus-level composition of the eight CES.** The communities' genus level composition as a function of dilution rounds are plotted. In cases where the genus is not assigned, the next higher assigned taxonomic rank is used in the label with: Fam - family, Cls - Class, Ord -Order, Phy - Phylum. For such genera, the ESV label is also indicated (Supplemetary Data 1). Only those genera that have a relative abundance of 5% or more in at least one CES are included here. See Supplementary Data 2 for phylogenetic information of each Genus.

| Reagent | Volume |
|---|---|
| PCR grade water | 13 $\mu$L |
| Forward primer (10 $\mu$M) | 0.5 $\mu$L |
| Reverse primer (10 $\mu$M) | 0.5 $\mu$L |
| Template DNA | 1 $\mu$L |
| PCR Master Mix (2X) | 10 $\mu$L |

**Table S1:** Reagents for PCR

| Temperature | Time | Repeat |
|---|---|---|
| 94 C | 3 min | |
| 94 C | 45 s | 45x |
| 50 C | 60 s | 45x |
| 72 C | 90 C | 45x |
| 72 C | 10 min | |
| 4 C | hold | |

**Table S2:** Thermocycler settings

| Parameter | Value | Unit | Definition | Source |
|---|---|---|---|---|
| $V_g$ | 0.02 | $L$ | Vial gas volume | - |
| $V_l$ | 0.02 | $L$ | Vial liquid volume | - |
| $T$ | 30 | °C | Temperature | - |
| $pH$ | 6.5 | 1 | pH | - |
| $R$ | 0.08205<br>$8.314 \times 10^{-3}$ | $L \cdot \text{atm} \cdot \text{mol}^{-1} \cdot K^{-1}$<br>$kJ \cdot \text{mol}^{-1} \cdot K^{-1}$ | Gas constant | [50] |
| $H_{O_2}$ | $1.27 \times 10^{-3}$ | $\text{mol} \cdot L^{-1} \cdot \text{atm}^{-1}$ | $O_2$ Henry's law constant (at $298.15K$) | [47] |
| $H_{CO_2}$ | $3.44 \times 10^{-2}$ | $\text{mol} \cdot L^{-1} \cdot \text{atm}^{-1}$ | $CO_2$ Henry's law constant (at $298.15K$) | [47] |
| $A_{O_2}$ | $-161.6$ | 1 | Parameter for $H_{O_2}$ | [47] |
| $B_{O_2}$ | 8160 | $K$ | Parameter for $H_{O_2}$ | [47] |
| $C_{O_2}$ | 22.39 | 1 | Parameter for $H_{O_2}$ | [47] |
| $A_{CO_2}$ | $-123.3$ | 1 | Parameter for $H_{CO_2}$ | [47] |
| $B_{CO_2}$ | 7335 | $K$ | Parameter for $H_{CO_2}$ | [47] |
| $C_{CO_2}$ | 16.739 | 1 | Parameter for $H_{CO_2}$ | [47] |
| $pK_a$ | 6.351 | 1 | $pK_a = -\log_{10} k_a$ for S8 (at $298.15K$) | [50] |
| $pK_2$ | 10.329 | 1 | $pK_2 = -\log_{10} k_2$ for S9 (at $298.15K$) | [50] |
| $\Delta_r H_a^\circ$ | 9.15 | $kJ \cdot \text{mol}^{-1}$ | Standard enthalpy of reaction for S8 | [50] |
| $\Delta_r C_{p\,a}^\circ$ | $-0.371$ | $kJ \cdot \text{mol}^{-1} \cdot K^{-1}$ | Standard heat capacity of reaction for S8 | [50] |
| $\Delta_r H_2^\circ$ | 14.70 | $kJ \cdot \text{mol}^{-1}$ | Standard enthalpy of reaction for S9 | [50] |
| $\Delta_r C_{p\,2}^\circ$ | $-0.249$ | $kJ \cdot \text{mol}^{-1} \cdot K^{-1}$ | Standard heat capacity of reaction for S9 | [50] |

**Table S3:** Pressure conversion parameters [50].

| Compound | Concentration |
|---|---|
| $C_6H_{12}O_6$ (glucose) | 1.666 mM |
| $NH_4Cl$ | 8 mM |
| $KH_2PO_4$ | 2.1 mM |
| $K_2HPO_4$ | 2 mM |
| $MgSO_4$ | 0.1 mM |
| $CaCl_2$ | 1 mM |
| $C_{10}H_{16}N_2O_8$ (EDTA) | 5.5 $\mu$M |
| $FeSO_4$ | 5.5 $\mu$M |
| $H_3BO_4$ | 15 $\mu$M |
| $ZnSO_4$ | 0.5 $\mu$M |
| $MnCl_2$ | 3.5 $\mu$M |
| $Na_2MoO_4$ | 0.58 $\mu$M |
| $CuSO_4$ | 0.15 $\mu$M |
| $Co(NO_3)_2$ | 0.8 $\mu$M |
| NaOH | 999 $\mu$M |
| $FeSO_4 \cdot 7H_2O$ | 999 $\mu$M |
| NaCl | 999 $\mu$M |

**Table S4:** Modified 1/2x Taub medium composition

| Nutrient | Source | Conc. (atoms) | Moles (atoms) | Mass [g] |
|---|---|---|---|---|
| Carbon | Glucose | 10mM | $2 \times 10^{-4}$ | $2.4 \times 10^{-3}$ |
| Nitrogen | Ammonia | 8mM | $1.6 \times 10^{-4}$ | $2.2 \times 10^{-3}$ |
| Phosphorous | Phosphate | 4mM | $8 \times 10^{-5}$ | $2.4 \times 10^{-3}$ |
| Atmosphere | | Conc. | | |
| $O_2$ (g) | Air | 21% | $1.8 \times 10^{-4}$ | $5.7 \times 10^{-3}$ |
| $CO_2$ (g) | Air | 0.4% | $3.5 \times 10^{-6}$ | $1.5 \times 10^{-4}$ |
| $O_2$ (l) | Dissolved | 0.27mM | $5.46 \times 10^{-6}$ | $1.7 \times 10^{-4}$ |
| $CO_2$ (l) | Dissolved | 0.12mM | $2.56 \times 10^{-5}$ | $1.1 \times 10^{-3}$ |

**Table S5:** Initial quantities of nutrients

| Compound excreted by *C. reinhardtii* | Corresponding compound in the Ecoplate | Mean ± Standard deviation [1/h] of consumption rate | |
|---|---|---|---|
| | | Round 1 | Round 4 |
| 2-O-Glycerol-$\alpha$-d-galactopyranoside | | | |
| Digalactosylglycerol | | | |
| Erythritol | i-Erythritol | 0.012 ± 0.002 | 0.012 ± 0.000 |
| Galacotse | | | |
| Glyceric acid | | | |
| Inositol,myo | | | |
| Malic acid | D-Malic acid | 0.019 ± 0.007 | 0.023 ± 0.008 |
| Nicotinamide | | | |
| Proline | | | |
| Putrescine | Putrescine | 0.025 ± 0.009 | 0.036 ± 0.018 |
| Pyroglutamic acid | | | |
| Ribose | | | |
| Threitol | | | |
| Threonic acid | L-Threonine | 0.012 ± 0.006 | 0.013 ± 0.003 |

**Table S6:** Compounds excreted in significant amounts by *Chlamydomonas reinhardtii* grown on its own
